# A prophage-encoded anti-phage defense system that prevents phage DNA packaging by targeting the terminase complex

**DOI:** 10.1101/2025.02.27.640495

**Authors:** Gil Azulay, Nadejda Sigal, Keren Tzohar Rabinovich, Dan Koenigsfeld, Olga Stadnyuk, Avijit Das, Polina Lisitzin, Einat Biderman, Ilya Borovok, Anat A. Herskovits

**Affiliations:** The Shmunis School of Biomedicine and Cancer Research, The George S. Wise Life Sciences Faculty, Tel Aviv University, Ramat Aviv, Tel Aviv 69978, Israel

**Keywords:** anti-phage defense, *Listeria monocytogenes*, phage DNA packaging, self-immunity, terminase

## Abstract

A unique feature of temperate phages is the ability to protect their host bacteria from a second phage infection. Such protection is granted at the lysogenic state, where the phages persist as prophages integrated within the bacterial chromosome, expressing genes that defend the host and themselves from predation. Here, we report a prophage-encoded anti-phage defense system that inhibits DNA packaging of invading phages in *Listeria monocytogenes*. This system includes a defense protein, TerI, and two self-immunity proteins, anti-TerI1 and anti-TerI2. TerI targets the terminase complex of invading phages to prevent DNA translocation into procapsids without halting the lytic cycle, leading to the release of unpacked non-infectious procapsids upon bacterial lysis. In contrast, the self-immunity proteins, anti-TerI1 and anti-TerI2, counteract TerI during prophage induction to allow virion production. This unique prophage-encoded anti-phage defense system, TERi, is prevalent in *Listeria* phages, providing population-level host protection without compromising the prophage lytic lifecycle.

## Introduction

Bacteriophages (or phages) are viruses that exploit bacterial cells for propagation, which are classified as lytic or temperate, based on their reproduction lifecycle.^1^ While lytic phages immediately enter a lytic cycle, in which they produce progeny virions that are released into the environment via bacterial lysis, temperate phages can integrate their genome into the bacterial chromosome and persist as prophages that replicate together with the host, a physiological state termed lysogeny.^2,3^ Under stress conditions, these prophages can switch into the lytic cycle by excising their genome from the bacterial chromosome and expressing genes that lead to the production and release of progeny virions, a process referred to as prophage induction.^1^ Despite this lytic ability of prophages, lysogeny is considered a long-term bacteria-phage relationship, which is maintained at the population level. During lysogeny, the fitness interests of the bacteria and the phage co-align, leading to the development of mutually beneficial interactions that support their co-existence under certain circumstances.^3–6^ An example of such interactions is the ability of prophages to protect their host, and hence themselves, from predation by exogenous phages. This type of protection was shown to be mediated by diverse mechanisms, some of which directly protect infected cells, while others provide protection at the population level. For instance, superinfection exclusion (Sie) systems prevent phage adsorption or DNA injection and thus protect the cells, whereas abortive infection (Abi) systems trigger programmed cell death or dormancy upon infection, thus sacrificing infected cells to avoid phage production and spread within the population.^7–10^

Our lab previously uncovered an unusual cooperative interaction between *Listeria monocytogenes* (*Lm*) strain 10403S and its sole prophage ϕ10403S that supports the survival of *Lm* during mammalian cell infection.^11–13^ *Lm* is an intracellular bacterial pathogen and the causative agent of Listeriosis disease in humans.^14,15^ It invades a wide array of mammalian cells, including gut, brain, and immune cells.^16^ Upon invasion, *Lm* is found within vacuoles or phagosomes (in the case of phagocytic cells), from where it escapes into the host cell cytosol to replicate.^17–19^ It has long been known that *Lm* strains carry prophages within their genome, yet the impact of this phenomenon on bacterial virulence was unclear. ϕ10403S belongs to a large group of *Listeria* phages of the *Siphoviridae* family that are integrated within the *comK* gene (*comK*-prophages), thus rendering this gene inactive.^20,21^ That said, our lab discovered that ComK plays a role in the escape of *Lm* from macrophage phagosomes into the cytosol. ComK expression in the macrophage phagosomes was found to rely on the excision of the prophage, yet unlike classic prophage excision (i.e., induction), this process did not lead to the production of progeny virions or bacterial lysis in the intracellular environment.^11–13^ Instead, the phage DNA was found to be maintained as an episome that does not express lytic genes (specifically, late genes), thereby allowing ComK expression and *Lm* phagosomal escape without triggering virion production and bacterial lysis. Moreover, the phage DNA was observed to re-integrate into *comK* during *Lm* intracellular growth, shutting off *comK* expression while resuming the lysogenic state.^11^ These findings revealed a cooperative phage behavior specific to the mammalian niche in which the prophage acts as an intervening DNA element that regulates bacterial gene expression to support the virulence of its host, a behavior termed *active lysogeny*.^4,22^

In this study, we report an additional symbiotic interaction between ϕ10403S and its host, which involves a unique phage-encoded anti-phage defense system that protects the *Lm* population from predation by exogenous phages. We found that ϕ10403S prophage encodes a small defense protein that targets the terminase complex of invading phages, thereby inhibiting their DNA packaging and virion assembly, as well as their epidemic spread within the host population. Furthermore, we found that the prophage encodes two self-immunity proteins that neutralize the defense protein during lytic induction, allowing for proper assembly of ϕ10403S virions and their release into the environment. As such, this study characterizes a novel prophage-encoded anti-phage defense system prevalent in *Lm* prophages that targets DNA packaging of invading phages without compromising the lytic lifestyle of the prophage.

## Results

### ϕ10403S encodes a small protein that inhibits virion production

Transcriptome analysis of ϕ10403S previously performed in our lab indicated a gene located at the negative strand of the phage early lytic gene module (opposite to *LMRG_02984*, **Figures 1A** and **S1**) that is constitutively transcribed under lysogenic and lytic conditions.^11^ The transcription profile of this gene was somewhat intriguing, as it is one of the few phage genes transcribed during lysogeny and the only one located within the early lytic gene module.^11^ It comprises an open reading frame (ORF) of 65 amino acids, hence named hereafter *orf65*, which overlapped the ORF of the *LMRG_02984* gene on the opposite strand (**Figure 1A**). We found *orf65* and its complementary gene (*LMRG_02984*), as well as their flanking genes *LMRG_01518* and *LMRG_02920,* to distribute among different *Lm* prophages with high conservation (e.g., the *comK*, A006 and A500 prophages, **Figure S2A**), raising the hypothesis that the products of these genes may functionally associate (**Figures 1A** and **S2B-D**). These genes had no annotation or predicted function, except for *LMRG_02920,* which our lab previously showed to encode a dual-function phage regulator (named AriS) that controls the SOS response.^23^

**Figure 1.**
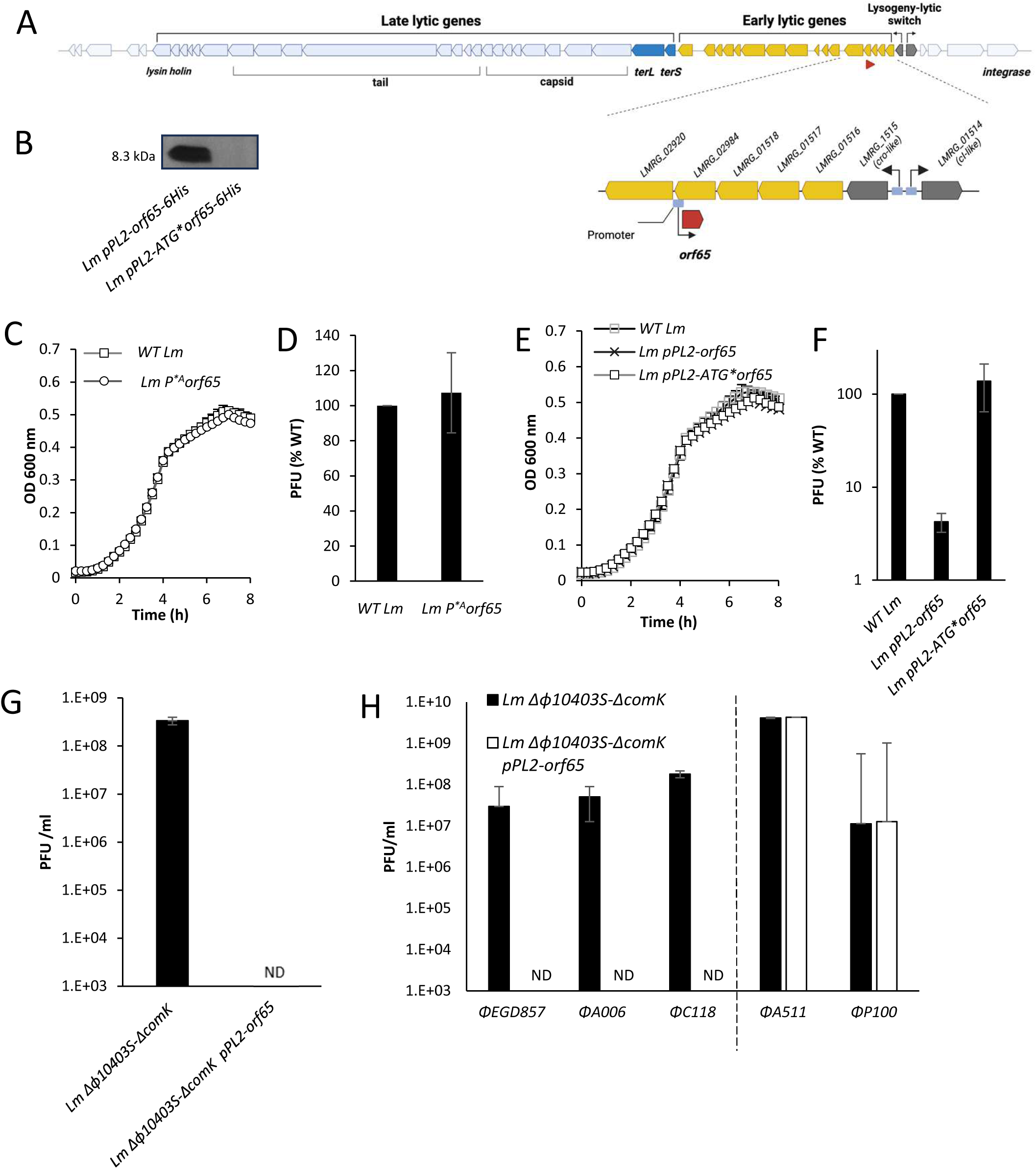
*orf65* encodes a protein that inhibits virion production upon prophage induction and exogenous phage infection. **A.** Schematic representation of the ϕ10403S genome and *orf65* genomic locus. **B.** Western blot analysis of ORF65 in bacteria producing His_6_-tagged ORF65 from the integrative plasmid pPL2 under the regulation of the constitutive *rpsD* promoter (*Lm pPL2-orf65-6His*), as compared to bacteria harboring an *orf65* clone mutated at the ATG start codon (*Lm pPL2-ATG*orf65-6His).* Bacteria were grown to exponential phase in BHI medium at 37°C, protein contents were extracted, and equal amounts of total protein extracts were analyzed. **C.** Growth analysis of WT *Lm* and *Lm* bacteria harboring a mutation in the promoter of *orf65* (*Lm P^*A^orf65*) in BHI medium at 30°C. The experiment was performed three times. Error bars represent the standard deviation of independent experiments. **D.** A plaque-forming assay of virions obtained from WT *Lm* and *Lm P^*A^orf65* bacteria upon MC treatment. Virions were isolated from bacterial lysates 6 h post-MC treatment and tested on the Mack861 indicator strain for plaque-forming units (PFU). Error bars represent the standard deviation of three independent experiments. **E.** Growth analysis of WT *Lm, Lm pPL2-orf65* and *Lm pPL2-ATG*orf65* strains in BHI medium at 30°C. The experiment was performed three times, and the error bars represent the standard deviation of independent experiments. **F.** A plaque-forming assay of virions obtained from WT *Lm, Lm pPL2-orf65* and *Lm pPL2-ATG*orf65* strains treated with MC. Virions were isolated from bacterial lysates 6 h post-MC treatment and tested on the Mack861 indicator strain for PFU. Error bars represent the standard deviation of three independent experiments. **G.** A plaque-forming assay of virions obtained upon ϕ10403S exogenous infection of *Lm Δϕ10403S*-*ΔcomK* and *Lm Δϕ10403S*-*ΔcomK+pPL2-orf65* strains. Error bars represent the standard deviation of three independent experiments. **H.** A plaque-forming assay of virions obtained upon exogenous infection of *Lm Δϕ10403S*-*ΔcomK* and *Lm Δϕ10403S*-*ΔcomK+pPL2-orf65* strains with the ϕEGD857, ϕA006, ϕC118, ϕA511, and ϕP100 phages. Error bars represent the standard deviation of three independent experiments.

To investigate the function of *orf65,* we first examined whether it translates into a protein. For this purpose, we generated a clone of *orf65* that encodes a six-histidine tag at the C-terminus of the predicted product and expressed the construct from the integrative pPL2 plasmid, using the constitutive *rpsD* promoter (pPL2*-orf65-6his*). We also generated a clone mutated at the ATG start codon of *orf65* to deliberately abolish protein translation, if it indeed occurs (pPL2-ATG\**orf65*-6his). The two clones were introduced into *Lm* 10403S bacteria (WT *Lm*), and ORF65 expression was evaluated by Western blot analysis using anti-His tag antibodies. The results indicated that *orf65* produces a small protein of 8.3 kDa, which was absent in bacteria carrying ATG-mutated *orf65* (**Figure 1B**). To investigate whether ORF65 plays a role during lysogeny or lytic induction, we generated an *Lm* strain harboring a mutation in the *orf*65 promoter at the “-10” recognition site of σA (*P^*A^orf65* mutant; **Figure S2E**), which impairs *orf65* transcription without altering the ORF of the complementary gene. Strand-specific real-time quantitative PCR (RT-qPCR) analysis confirmed that the *P^*A^orf65* mutation impaired *orf*65 transcription in comparison to the wild-type (WT) strain (>10-fold) (**Figure S3**). This mutant was then used to examine the importance of ORF65 under lysogenic conditions (i.e., during bacterial growth in BHI medium at 37°C) and upon lytic induction (i.e., upon mitomycin C (MC) treatment at 30°C), by monitoring bacterial growth and virion production in *P^*A^orf65* and WT bacteria, respectively. The results indicated that ORF65 does not affect bacterial growth during lysogeny nor virion production upon MC treatment (as estimated using a plaque-forming assay) (**Figures 1C, 1D**).

We next examined the impact of *orf65* ectopic expression on the bacteria and phage under lysogenic and lytic conditions. For this, we again cloned *orf65* and its ATG* variant into pPL2 plasmid, this time, however, without the histidine tag-encoding sequence, and introduced the resulting plasmids into WT *Lm* (*Lm* pPL2*-orf65* and *Lm* pPL2*-ATG*orf65*). Notably, whereas *orf65* ectopic expression did not affect *Lm* growth under lysogenic conditions, it strongly affected the production of virions upon MC treatment (**Figures 1E, 1F**). Specifically, *orf65*-expressing bacteria yielded a significantly lower number of virions than WT bacteria or bacteria carrying the *ATG*orf65* mutation, indicating that ORF65 somehow interfered with the lytic cycle of the prophage (**Figure 1F**). To exclude the possibility that this phenotype is related to *LMRG_02984*, such as due to an RNA-antisense silencing effect caused by overly transcribed *orf65*, we compared the transcription levels of *LMRG_02984* in bacteria expressing and not expressing *orf65*; no significant difference was found (**Figure S4A**). We, furthermore, found that *LMRG_02984* itself is not required for virion production, as a mutant deleted of this gene produced virions like WT bacteria upon MC treatment (Δ*LMRG_02984* vs. WT *Lm*; **Figure S4B**). Moreover, ectopic expression of ORF65 also inhibited virion production in the Δ*LMRG_02984* mutant, supporting the premise that ORF65 activity is independent of its complementary gene (**Figure S4B**). Intrigued by these findings, we examined whether ORF65 can further inhibit virion production upon exogenous phage infection (i.e., without MC treatment). To this end, we infected *Lm* bacteria cured of ϕ10403S and deleted of the *comK* gene with free particles of ϕ10403S in the presence or absence of *orf65* ectopic expression (i.e., using *Lm Δϕ10403S*-*ΔcomK* with or without the pPL2-*orf65* plasmid). We then assessed virion production three hours post-infection using a plaque-forming assay. The results demonstrated that ORF65 could efficiently block virion production upon exogenous phage infection, as no single plaque was detected in *orf65*-expressing bacteria that were exogenously infected, compared to control cells (**Figure 1G**). Notably, this activity of ORF65 was not specific to ϕ10403S, as ORF65 could further inhibit infection by other listerial *Sipho* phages (temperate), such as EGD857, C118 and A006, but not *Myo* phages like A511 and P100, which are strictly lytic (**Figure 1H**).

### ORF65 inhibits phage DNA packaging into procapsids

To decipher the mechanism by which ORF65 inhibits virion production, we examined its effect on the different steps of the phage lytic cycle. Since ORF65 inhibited virion production upon both phage infection and induction, we surmised that it acts downstream of phage DNA injection or excision, targeting, for example, phage DNA replication, gene transcription, virion assembly or phage-mediated cell lysis, all processes that can result in a low production of virions if hindered. As such, we examined the impact of ORF65 on ϕ10403S virion production upon MC treatment. We tested phage DNA replication, gene transcription and cell lysis in bacteria expressing or not expressing *orf65* using WT *Lm* with or without pPL2*-orf65*, respectively. Phage DNA replication was assessed using qRT-PCR, measuring the copy numbers of the phage *attP* site, which is formed only upon excision of the phage DNA from the bacterial chromosome. We further quantified the copy numbers of the *attB* site in the bacterial genome as a control (representing the intact *comK* gene) to evaluate the frequency of prophage excision within the population. In addition, we compared the transcription levels of representative early and late lytic genes in bacteria expressing *orf65* or not to assess its impact on the transcription profile of the phage. The results indicated that ORF65 does not affect phage DNA replication (or excision) nor transcription of lytic genes (**Figures 2A, 2B**), suggesting that it targets downstream processes of the phage lytic cycle. We next examined whether ORF65 prevents phage-mediated bacterial lysis by comparing the growth of MC-treated bacteria expressing or not expressing *orf65*; no difference was observed (**Figure 2C**). However, since it was previously demonstrated that *Lm* strain 10403S carries another phage-derived element that triggers cell lysis upon MC treatment, namely, the monocin element that encodes phage tail-like bacteriocins (monocins),^12,24,25^ we repeated the experiment, this time using a strain deleted of monocin structural and lysis genes (i.e., the Δ*mon*-*struc-lys* mutant). As shown in **Figure 2D**, similar results were obtained using this mutant, supporting the conclusion that ORF65 does not inhibit phage-mediated bacterial lysis.

**Figure 2.**
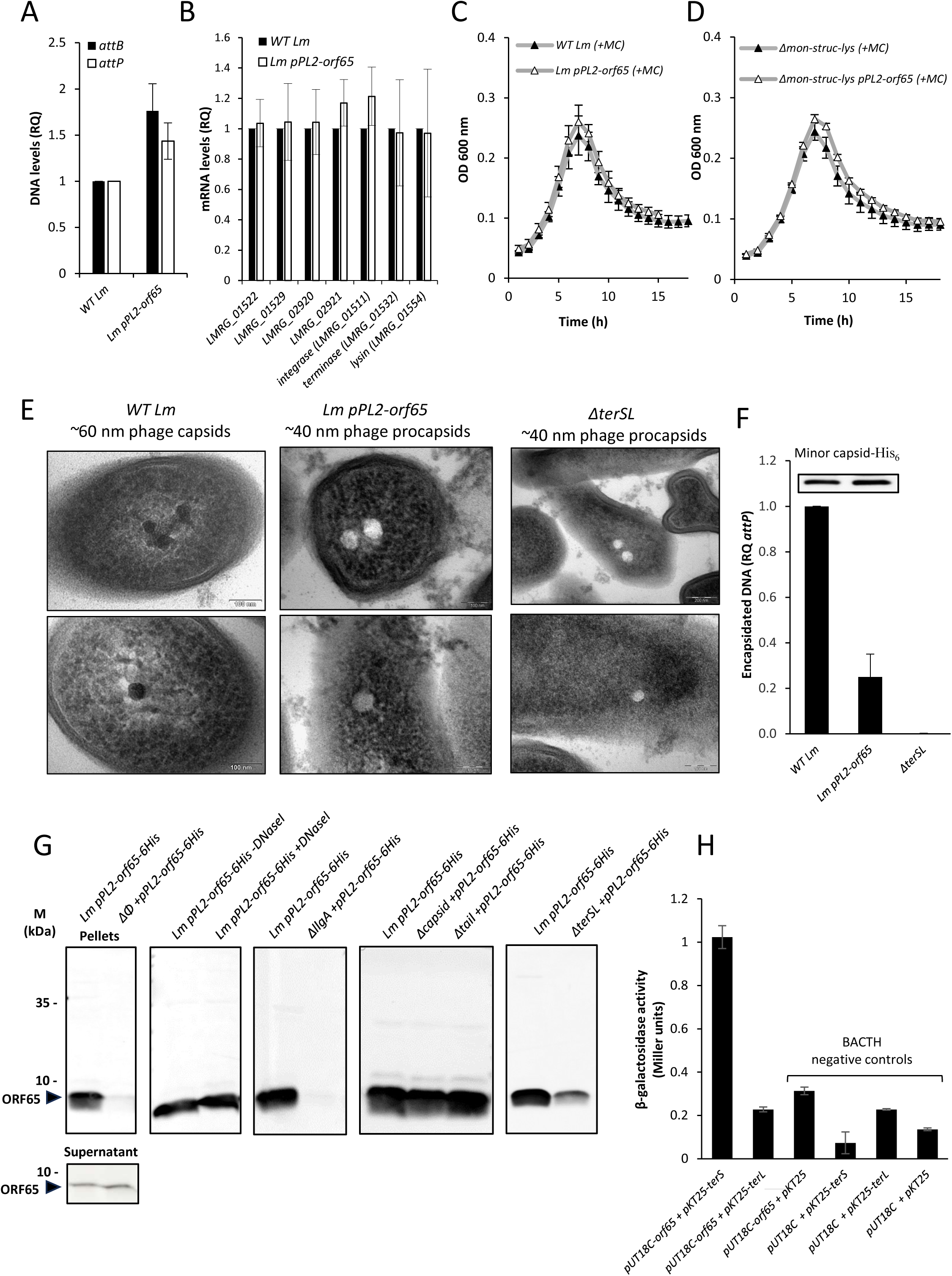
ORF65 blocks DNA packaging via interaction with the terminase complex. **A.** qRT-PCR analysis of ϕ10403S *attB* and *attP* sites (indicating phage excision and replication, respectively) upon MC treatment of WT *Lm* and *Lm pPL2-orf65* bacteria. The bacteria were grown in BHI medium with MC for 3 h. The data are presented as relative quantity (RQ), relative to *attB/P* levels in WT bacteria. The experiment was performed three times. Error bars represent the standard deviation of three independent experiments. **B.** Transcription analysis of representative early and late lytic genes of ϕ10403S in WT *Lm* and *Lm pPL2-orf65* bacteria using RT-qPCR. Indicated strains were grown in BHI medium with MC treatment for 3 h. The data are presented as RQ, relative to the corresponding levels in WT bacteria. The experiment was performed three times. Error bars represent the standard deviation of three independent experiments. **C-D.** Growth analysis of WT *Lm* (**C**) and a *Lm* mutant deleted of the structural and lysis genes of the monocin element (*Δmon-struc-lys)* (**D**) with or without *pPL2-orf65.* The strains were grown in BHI medium with MC (+MC) at 30°C. The experiment was performed three times. Error bars represent the standard deviation of three independent experiments. **E.** Transmission electron micrographs demonstrating ϕ10403S capsids (or procapsids) in WT *Lm*, in comparison to *Lm-pPL2-orf65* and *ΔterSL* strains taken 5 h post-UV irradiation at 30°C. Two representative images are shown for each strain. Experiments were performed three times. **F.** Analysis of encapsidated phage DNA using qRT-PCR analysis of the *attP* site in capsids/virions collected by ultra-centrifugation following DNaseI treatment from WT *Lm, Lm-pPL2-orf65* and *ΔterSL* bacterial cultures treated with MC. The experiment was performed three times. Error bars represent the standard deviation of three independent experiments. The upper panel shows Western blot analysis of a His_6_-tagged minor capsid protein (LMRG_01534) in capsids/virions samples collected from the corresponding strains for a control. The experiment was performed three times. The figure shows a representative experiment. Additional biological repeats can be found in the **Data S1 file**. **G.** A Western blot analysis of His_6_-tagged ORF65 in pellet samples containing ϕ10403S particles isolated from bacterial lysates of MC-treated bacteria by ultra-centrifugation. A supernatant control sample is also shown. *Lm* bacteria expressing His_6_-ORF65 were grown in BHI medium and treated with MC for 6 h. Bacterial lysates were filtrated, treated with DNaseI when indicated and subjected to ultra-centrifugation (100,000 g). The pellets were resuspended and analyzed. The experiment was done with different mutants as indicated to identify those phage components to which ORF65 binds. The experiment was performed three times. The figure shows a representative blot. Additional biological repeats can be found in **the Data S1 file**. **H.** A bacterial adenylate cyclase-based two-hybrid system was used to detect protein-protein interactions between ORF65 and TerS or TerL. *orf65* was cloned into the pUT18C plasmid, while the genes encoding for TerS and TerL were cloned into plasmid pKT25, with both plasmids provided with the BACTH kit. The plasmids were then co-transformed into *E. coli* strain BTH101. As negative controls, BTH101 bacteria were also transformed with plasmids pUT18C-*orf65* and pKT25, pUT18C and pKT25-*terS* or pKT25-*terL,* and pUT18C and pKT25. The strains were grown in LB medium supplemented with ampicillin, kanamycin and IPTG, and a β-galactosidase activity assay was performed using ONPG. β-galactosidase activity was normalized to the sample OD_600nm_ and is presented in Miller units.

Although the results indicated that the lytic cycle of the phage progresses appropriately, they did not provide information regarding the assembly of virions, i.e., whether virions are correctly assembled or not. To address this point, we used transmission electron microscopy (TEM) to examine virion production in bacteria expressing or not expressing *orf65*. Upon scanning dozens of stained bacterial sections, we observed that while WT bacteria exhibited heavily stained (dark) ∼60 nm capsids, bacteria expressing *orf65* exhibited smaller capsids of ∼40 nm that did not present internal staining, instead appearing as white, “ghost”-like capsids (**Figure 2E**). We hypothesized that these small ghost capsids are procapsids devoid of DNA, since DNA packaging is known to expand the capsid shell, which reaches its final size only upon full condensation of the phage genome.^26,27^ To test this hypothesis, we generated a mutant deleted of the phage terminase complex comprising the TerS and TerL subunits (Δ*terSL*), which is known to translocate phage DNA into procapsids.^26,27^ Notably, the Δ*terSL* mutant demonstrated ∼40 nm procapsids that did not present internal staining but rather appeared white, resembling the procapsids observed in *orf65*-expressing bacteria (**Figure 2E**). This finding supported the hypothesis that ORF65 interferes with phage DNA packaging, in turn inhibiting the production of progeny virions. To corroborate this premise, we examined whether the small procapsids released from ORF65-expressing bacteria lack DNA. For this purpose, we isolated capsids from bacterial lysates using high-speed centrifugation (100,000 g) and quantified encapsidated DNA by analyzing copy numbers of the phage *attP* site using qRT-PCR. Of note, this experiment was performed as a DNase I protection assay, in which only encapsidated DNA, which is protected from degradation, could be further detected by PCR following phenol extraction. The results of this experiment indicated that ORF65 prevents phage DNA packaging, as the amount of phage DNA recovered from capsids produced by *orf65*-expressing bacteria was significantly lower than that recovered from capsids produced by bacteria not expressing *orf65* (**Figure 2F**). At the same time, no DNA was detected in procapsids produced by the Δ*terSL* mutant (**Figure 2F**). As a control, we confirmed that both bacterial strains (i.e., strains expressing *orf65* or not) produced and released a similar number of capsids by detecting a representative capsid protein (LMRG_01534, a minor capsid protein) in a Western blot analysis using antibodies directed against the His-tagged version of this protein (**Figure 2F**).

To gain insight into the mechanism by which ORF65 inhibits phage DNA packaging, we considered whether it targets the phage DNA, the procapsid, or the terminase complex. For this purpose, we used the same high-speed centrifugal sedimentation approach as above in combination with Western blot analysis to determine whether ORF65 co-sediments with any of these complexes. Since ORF65 is a small protein, we reasoned it would not sediment upon centrifugation unless bound to a large macromolecule. To this end, bacteria expressing His-tagged ORF65 were grown in BHI medium and treated with MC, and their released phage particles (e.g., intact virions, procapsids and tails) were collected from cell lysates using high-speed centrifugation. The particle-containing pellets were then resuspended and subjected to Western blot analysis using anti-His antibodies to detect tagged ORF65. As shown in **Figure 2G**, ORF65 was detected in the pellet (as well as in the supernatant), with its sedimentation being absolutely dependent on the presence of the phage, as it was not detected in pellets prepared from phage-cured bacteria (*Δϕ10403S* + pPL2-*orf65-6his*). This finding confirmed that ORF65 binds a large macromolecule or complex of the phage. To assess whether ORF65 associates with phage DNA, we treated the bacterial lysates with DNase I before high-speed centrifugation. We found that ORF65 sedimentation was independent of phage DNA, as it was similarly detected in pellets prepared from lysates treated with DNase I or not (**Figure 2G**). To assess whether ORF65 interacts with one of the phage structural complexes, i.e., the capsid, the tail or the terminase, we repeated the experiment using a strain that is deleted of the *llgA* gene (*ΔllgA*), which encodes the phage main activator of the late lytic genes (i.e., *terS*, *terL*, *capsid*, *tail* and *lysis* genes).^11^ We previously demonstrated that *ΔllgA* mutant allows phage DNA excision and replication yet lacks the expression of the structural and lysis genes. Notably, ORF65 was not detected in pellets prepared from the Δ*llgA* mutant, supporting the hypothesis that ORF65 is associated with one of the late structural complexes (i.e., the terminase, the capsid or the tail) (**Figure 2G**). To determine which complex binds ORF65, we examined mutants deleted of all genes encoding each complex, named here *Δcapsid, Δtail* and *ΔterSL* (**Data S1 File**). Interestingly, the results indicated ORF65 sedimentation to be independent of the phage capsid or tail but largely dependent on the presence of the terminase (**Figure 2G**). To examine whether ORF65 directly binds the terminase complex, we used the bacterial adenylate cyclase-based two-hybrid (BACTH) system to monitor protein-protein interactions between ORF65 and each of the terminase subunits, i.e., the small subunit TerS and the large subunit TerL. The results indicated that ORF65 directly binds TerS (**Figure 2H**), supporting the hypothesis that ORF65 inhibits phage DNA packaging via direct interaction with the terminase complex. Since we observed that ORF65 sedimentation is independent of the presence of capsids (**Figure 2G**), we concluded that ORF65 binds the terminase complex in its free form before associating with the pre-assembled procapsid, possibly preventing their interaction and hence, DNA packaging. Considering these findings, we named ORF65 TerI, reflecting its newly described function as a terminase inhibitor.

### ϕ10403S encodes two self-immunity proteins that counteract TerI activity

Having discovered that ϕ10403S encodes a protein that inhibits phage DNA packaging, we surmised that it acts as an anti-phage defense protein during lysogeny, preventing virion assembly by invading phages and thus their spread within the host population. That said, the data indicated that *terI* is also transcribed upon lytic induction of ϕ10403S (**Figure S1**), implying that the prophage must have a counter-defense system that counteract TerI activity to allow virion production. In a search for such a system, we screened for suppressor mutations in the ϕ10403S genome that rescue virion production upon ectopic expression of *terI*. For this purpose, we performed successive cycles of ϕ10403S infections of phage-cured *terI*-expressing bacteria (Δ*ϕ10403S*-pPL2-*terI*) to enrich for phage mutants that produce a high titer of virions, despite the presence of TerI. In this experiment, we used a ϕ10403S-phage that carries a kanamycin resistance gene (ϕ10403S-*kan^R^*) to allow for the selection of lysogens in each infection cycle.^11^ Bacteria were infected with ϕ10403S-*kan^R^*phage particles and selected for lysogens. The lysogens were then subjected to UV irradiation to trigger phage induction, and the resulting virions were collected, quantified in a plaque-forming assay and used to initiate another round of phage infection of *terI*-expressing phage-cured bacteria. Upon performing this experiment for multiple cycles (n=10) and using independent lines (n=4), we detected one line that demonstrated an increase in virion production, as compared to a control experiment performed with a fresh WT phage in each cycle. Sequencing the genomes of 17 independent lysogens recovered from this line, we found them all to carry the same mutation, namely, a single nucleotide substitution (T to C) in the lysogenic-lytic regulatory region immediately upstream of the *cro*-like gene (*LMRG_01515*, like the *cro* of λ-phage) (**Figure 3A**, upper panel and **Figure S5**). As shown in **Figure 3A**, this mutation provided full immunity against TerI, as the mutated phage (ϕ*-suppressor*) produced an equal number of virions in bacteria producing the protein or not (**Figure 3A**). Importantly, virion production by WT bacteria carrying the mutated phage was similar to that of WT bacteria carrying a WT phage (not expressing *terI*), indicating that the mutation has no general enhancing effect on virion production but rather specifically rescues virion production in the presence of TerI (**Figure 3A**). Based on the location of the mutation (and information from the λ-phage), we predicted that the mutation targeted the operator site of Cro and hence affected its binding (i.e., lowered its affinity), thus resulting in increased expression of the phage early lytic genes. To examine this hypothesis, we tested the binding of Cro (LMRG_01515) to its putative operator site by performing an electromobility shift assay (EMSA), using increasing amounts of purified recombinant His_6_-tagged Cro (Cro-6His) and probes containing the operator site with or without the suppressor T to C mutation (**Figure S5**). In line with our hypothesis, the EMSA analysis indicated that Cro binds both probes, albeit with different affinities, demonstrating a K_D_ of 321 nM for the probe containing the native site and 433 nM for the probe containing the mutated site (**Figure S5**). Next, we analyzed the transcription level of *cro* (corresponding to the first early gene) upon induction of ϕ10403S and ϕ*-suppressor* in bacteria expressing *terI* or not. The results indicated that irrespective of *terI* expression, the mutated phage (ϕ*-suppressor*) exhibited augmented transcription of the early genes (> 3-fold), as compared to the WT phage (**Figure 3B**), raising the hypothesis that one of the early genes encodes a factor that counteracts the activity of TerI. To test this possibility, we ectopically expressed each of the first early genes, in addition to *terI*, and searched for gene candidates that rescue virion production in the presence of TerI (i.e., producing virions like WT bacteria not expressing TerI). For this purpose, we used the integrative plasmids pPL2 and pSNIB1, the latter having been specifically designed for this study (see Material and methods). To this end, the *terI* gene was cloned in the pSNIB1 plasmid, and each of the early genes, i.e., *LMRG_01515, LMRG_01516*, *LMRG_01517*, *LMRG_01518* and *LMRG_*02984, were cloned into plasmid pPL2. We next evaluated virion production in bacteria carrying these plasmids and found two gene candidates, *LMRG_01518* and *LMRG_02984*, whose ectopic expression rescued virion production despite the presence of TerI (**Figure 3C**). As described earlier, *LMRG_02984* is the complementary gene of *terI*, and *LMRG_01518* is one of their flanking genes (**Figure 3A**). Ectopic expression of each of these genes alone (i.e., without *terI* expression) did not affect virion production, as compared to WT bacteria, suggesting that the encoded proteins play a specific role in neutralizing TerI (**Figure 3D**). That said, when performing this experiment, we noticed that co-expression of *terI* and *LMRG_02984* resulted in decreased *terI* transcription, likely due to an anti-sense RNA silencing effect, which was not observed with *LMRG_01518* (**Figure S6A**). In light of this possibility, we repeated the experiment, this time using a synthetic *terI* gene that contained codons that were altered without changing the encoded amino acid sequence (pSNIB1-*syn-terI*) so as to prevent possible annealing to *LMRG_02984* transcripts. Initially, we assessed whether the synthetic *terI* gene produced a functional protein that inhibits virion production when over-expressed and then co-expressed it with *LMRG_02984*. The results demonstrated that the synthetic gene produced a fully active protein and that LMRG_02984 could rescue its phenotype, i.e., alleviating virion production in the presence of TerI as in WT bacteria (this time without the silencing effect) (**Figures 3E** and **S6B**). Similar results were obtained with *LMRG_01518* and the synthetic *terI* gene (**Figure 3E**).

**Figure 3.**
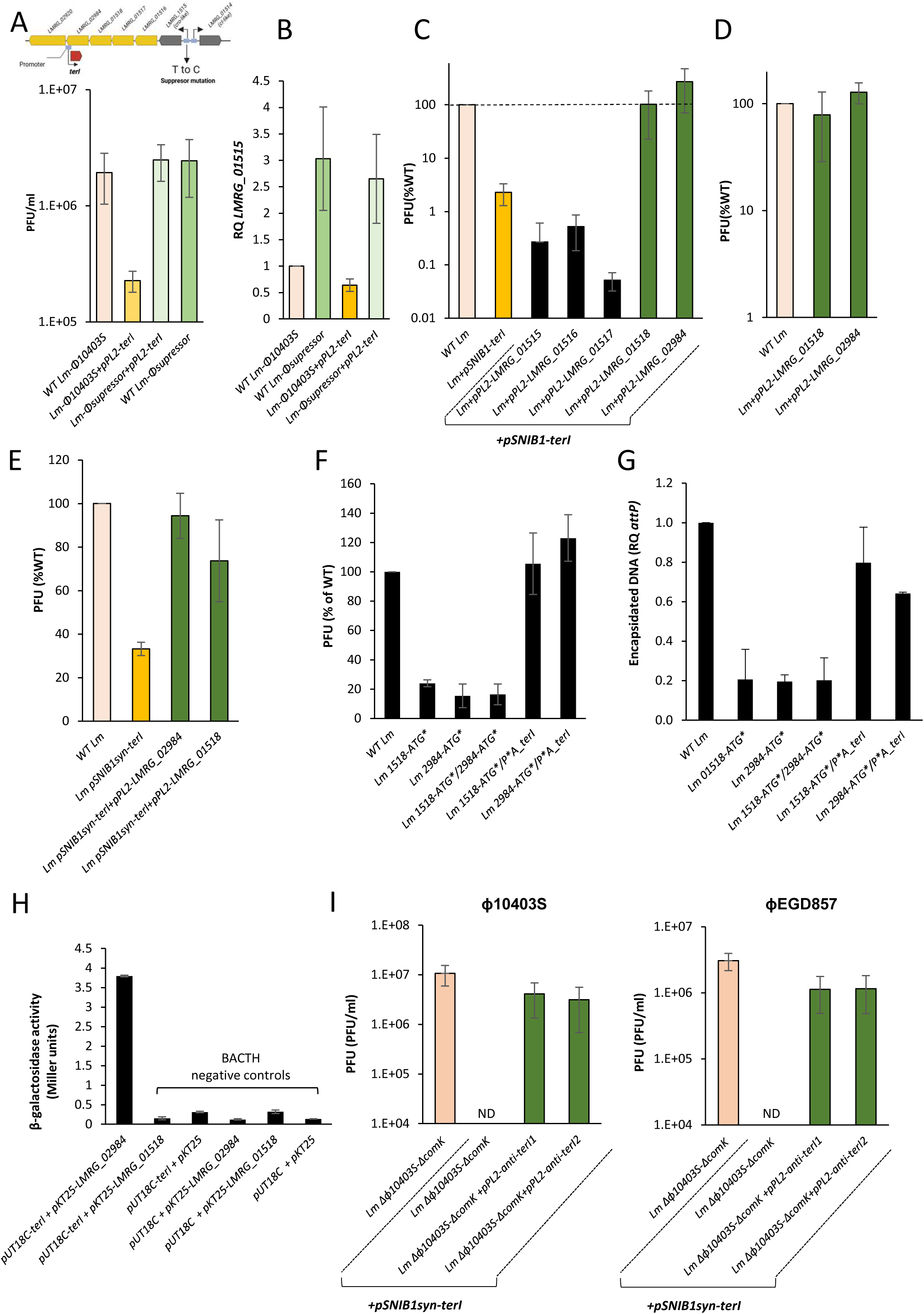
ϕ10403S phage encodes two self-immunity proteins that counteract TerI. **A.** A plaque-forming assay of virions obtained from WT *Lm* and *Lm ϕ-suppressor* strains upon MC treatment, using bacteria expressing TerI from plasmid pPL2 (*pPL2-terI*) or not. Virions were filtrated from bacterial lysates (6 h post-MC treatment) and tested on the Mack861 indicator strain for PFU. Error bars represent the standard deviation of three independent experiments. **B.** RT-qPCR analysis of the transcriptional levels of *LMRG_01515* gene in WT *Lm* and *Lm ϕ-suppressor* strain expressing TerI (from plasmid *pPL2-terI*) or not. Indicated strains were grown in BHI medium with MC treatment for 1 h. The results are presented as RQ, relative to the levels in WT bacteria. Error bars represent the standard deviation of three independent experiments. **C.** A plaque-forming assay of virions obtained from WT *Lm* expressing the indicated early lytic genes from the pPL2 plasmid using a TetR-dependent promoter and the *terI* gene from the pSNIB1 plasmid (*pSNIB1-terI*) under MC treatment. Bacterial cultures were grown in the presence of the *tet*-promoter inducer, anhydrotetracycline (AT, 100 ng/ml) and MC. Virions were obtained 6 h post-MC treatment and tested on the Mack861 indicator strain for PFU. Error bars represent the standard deviation of three independent experiments. **D.** Plaque-forming assay of virions obtained from WT *Lm* and *Lm* bacteria expressing *LMRG_01518 or LMRG_02984* from the pPL2 plasmid upon MC treatment. Bacterial cultures were grown in the presence of the *tet*-promoter inducer anhydrotetracycline (AT, 100 ng/ml) and MC. Virions were tested 6 h post-MC treatment using the Mack861 indicator strain for PFU. Error bars represent the standard deviation of three independent experiments. **E.** Plaque-forming assay of virions obtained from WT *Lm* and *Lm* bacteria expressing a synthetic clone of *terI,* harboring alternative codons (*pSNIB1-syn-terI*) and *LMRG_01518* or *LMRG_02984* from the pPL2 plasmid upon MC treatment. Bacterial cultures were grown in the presence of the *tet*-promoter inducer, anhydrotetracycline (AT, 100 ng/ml) and MC. Virions were tested 6 h post-MC treatment using the Mack861 indicator strain for PFU. Error bars represent the standard deviation of three independent experiments. **F.** A plaque-forming assay of virions obtained from WT *Lm* and a *Lm* strain mutated at the first methionine codon of *LMRG_01518* (*Lm 1518-ATG\**) or *LMRG_02984* (*Lm 2984-ATG\**) or both (*Lm 1518-ATG*/2984-ATG\**) upon MC treatment. Virions were tested 6 h post-MC treatment using the Mack861 indicator strain for PFU. Error bars represent the standard deviation of three independent experiments. **G.** Analysis of encapsidated DNA using qRT-PCR analysis of the *attP* site in capsids/virions collected by ultracentrifugation following DNaseI treatment from WT *Lm* and indicated strains upon MC treatment. The experiment was performed three times. Error bars represent the standard deviation of three independent experiments. **H.** A bacterial adenylate cyclase-based two-hybrid system was used to detect protein-protein interactions between TerI and LMRG_01518 or LMRG_02984. *terI* was cloned into the pUT18C plasmid, while the genes encoding for LMRG_01518 and LMRG_02984 were cloned into plasmids pKT25, with both plasmids provided with the BACTH kit. The plasmids were then co-transformed into *E. coli* strain BTH101. As negative controls, BTH101 bacteria were also transformed with plasmids pUT18C-*terI* and pKT25, pUT18C and pKT25-*LMRG_01518* or pKT25-*LMRG_01518,* and pUT18C and pKT25. The strains were grown in LB medium supplemented with ampicillin, kanamycin and IPTG, and β-galactosidase activity assay was performed using ONPG. β-galactosidase activity was normalized to the sample OD_600nm_ and is presented in Miller units. **I.** A plaque-forming assay of ϕ10403S (left panel) and ϕEGD857 (right panel) virions obtained upon exogenous infection of *Lm Δϕ10403S*-*ΔcomK* and *Δϕ10403S-ΔcomK+pSNIB1-terI* strains expressing *LMRG_01518* or *LMRG_02984* from the pPL2 plasmid or not. Error bars represent the standard deviation of three independent experiments.

Since the data revealed that LMRG_01518 and LMRG_02984 act as factors that counteract the activity of TerI, we next examined their *in vivo* role during ϕ10403S induction (i.e., upon MC treatment). Accordingly, we generated mutants that do not express these proteins by introducing mutations at the ATG start codons of their encoding genes, either separately or together (i.e., the *Lm 1518-ATG\** and *Lm 2984-ATG\** single mutants and the *Lm 1518-ATG*/2984-ATG\** double mutant). We next tested the mutant bacteria for virion production upon MC treatment and found them all to be severely impaired, producing only ∼20% of the virions, as compared to WT *Lm* carrying a WT phage (**Figure 3F**). This phenotype was completely dependent on the expression of *terI*, as combining the above mutations with the *P*^A^terI* mutation (which impairs *terI* transcription) resulted in a complete rescue of virion production (**Figure 3F**). In line with TerI activity as inhibitor of phage DNA packaging, we found that the *1518-ATG*\* and *2984-ATG*\* mutants, as well as the *1518-ATG*\*/*2984-ATG*\* double mutant, produced and released (pro)capsids devoid of DNA (**Figure 3G**). This phenotype was also dependent on *terI* expression, as it disappeared when the *P*^A^terI* mutation was introduced, demonstrating for the first time the endogenous activity of TerI in the course of the lytic cycle (**Figure 3G**). We next examined whether LMRG_01518 and LMRG_02984 interact with TerI using the BACTH system, which detected direct interaction between TerI and LMRG_02984 (**Figure 3H**).

Altogether, these findings indicated that TerI is fully active during ϕ10403S induction and that LMRG_01518 and LMRG_02984 counteract its activity, functioning as self-immunity proteins that allow ϕ10403S DNA packaging and virion assembly. We, therefore, named LMRG_01518 and LMRG_02984, anti-TerI1 and anti-TerI2, respectively. Finally, we asked whether these proteins can further rescue virion production upon exogenous phage infection of *terI*-expressing bacteria. To address this question, two phages were tested, ϕ10403S and ϕEGD857, infecting *Lm Δϕ10403S*-*ΔcomK* bacteria expressing TerI and each of the anti-TerI proteins. The results demonstrated that each self-immunity protein could neutralize TerI anti-phage activity when over-expressed, leading to the production and release of infective virions, as observed in bacteria not expressing the *terI* gene (**Figure 3I**). While *in vivo* experiments indicated that both proteins are required to neutralize TerI (**Figure 3 F-G**), the over-expression experiments showed that each protein alone can neutralize TerI, a discrepancy that might be explained by their level of expression (**Figures 3E, 3I**). Altogether, these experiments uncovered a novel prophage-encoded anti-phage defense system that inhibits DNA packaging, named TERi, comprising a defense protein and two self-immunity proteins that provide anti-phage protection at the population level without compromising the lytic lifestyle of the encoding phage.

### The TERi genomic locus contains different genes that encode TerI-like activity

Having discovered the TERi system, we wondered how widespread it is among *Lm comK*-prophages. Upon analyzing the genomes of 149 *comK*-prophages of different *Lm* strains, we found *terI* and its two self-immunity genes in 38% of the genomes. That said, we found the promoter region of *terI* to be highly conserved, i.e., present in 96% of the *comK*-prophages, exhibiting the same sequence, location and orientation as in ϕ10403S (**Figures 4A** and **S7**). Downstream to this promoter, we identified a set of four different genes that alternated with *terI*, namely, *orf80, orf67, orf180* and *orf69*, which were found in 30.9%, 14.8%, 6.7% and 5.4% of *comK*-prophages, respectively (**Figure 4A**). Although none of these genes shared sequence similarity with *terI,* two of them (*orf80* and *orf180*) were found to associate with *anti-terI1*, which was found nearby in 95.6% and 70% of the cases, respectively (**Figure 4A**). Considering that defense genes often cluster or locate at specific loci (hotspots),^28,29^ we tested the possibility that these genes also encode anti-phage defense proteins. We, therefore, cloned *orf80* (the second most abundant gene in this locus) and *orf180* (which appears much less frequently) into pPL2 plasmids and introduced these vectors into *Lm* strain 10403S (*Lm* pPL2*-orf80* and *Lm* pPL2*-orf180*). Next, we examined virion production in MC-treated bacteria ectopically expressing *orf80* or *orf180* and obtained similar results to those observed with *terI* expression. Ectopic expression of each protein efficiently inhibited ϕ10403S virion production, suggesting that ORF80 and ORF180 also possess anti-phage defense activity (**Figure 4B**). Similarly to TerI, these proteins did not affect ϕ10403S DNA replication or excision (**Figure 4C**) but instead inhibited phage DNA packaging, as reflected by the low amount of encapsidated DNA in procapsids produced in their presence (**Figure 4D**). Moreover, ectopic expression of *orf80* or *orf180* further inhibited virion production upon exogenous phage infection using ϕ10403S and ϕEGD857, supporting the conclusion that they function as anti-phage defense proteins that inhibit DNA packaging and, hence, virion assembly (**Figure 4E**). These proteins were, therefore, named TerI-80 and TerI-180. These findings thus revealed that targeting DNA packaging as an anti-phage defense is a common strategy of *Listeria* prophages, a strategy that has likely evolved multiple times independently.

**Figure 4.**
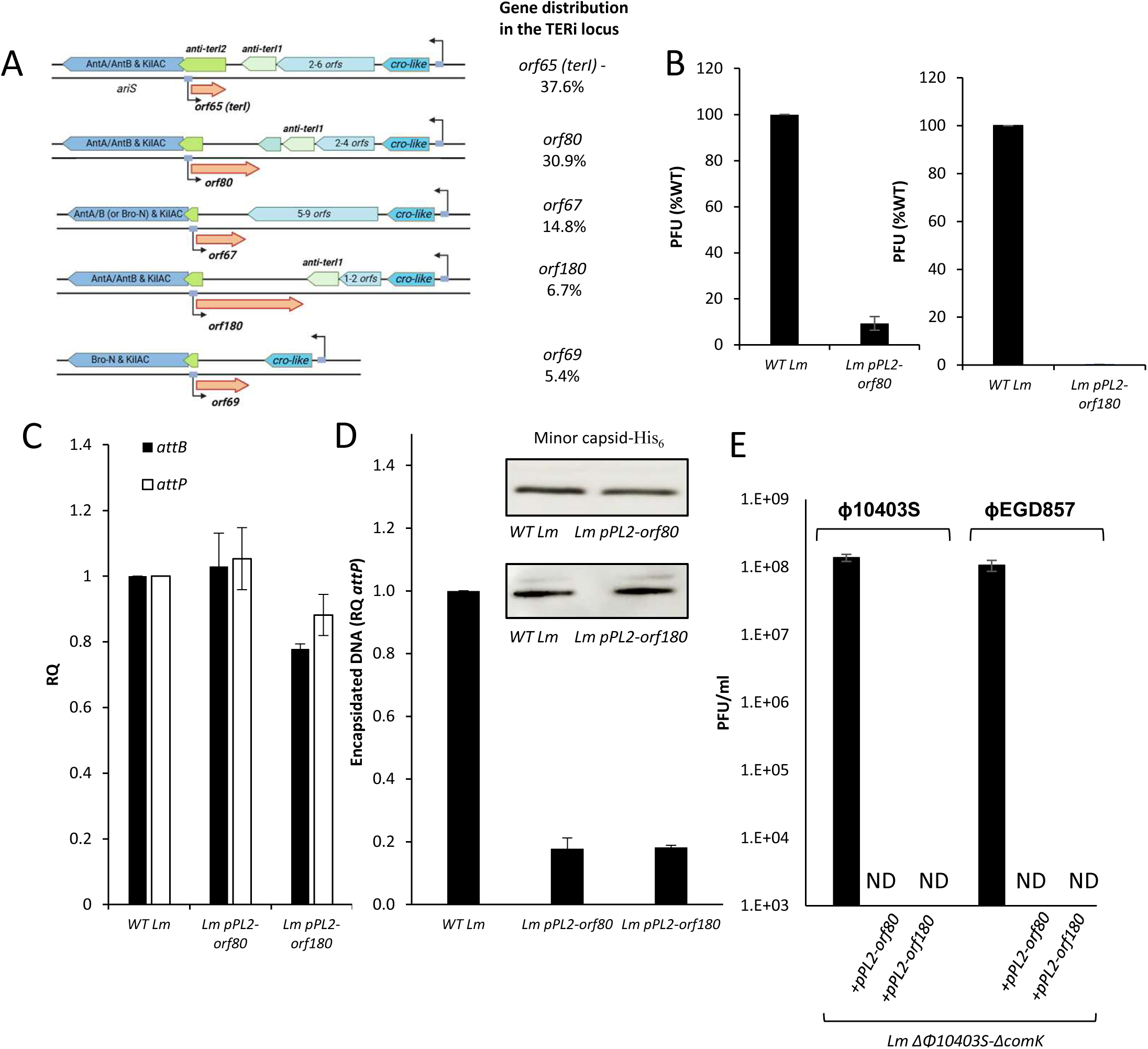
The *terI* gene locus is conserved, serving as a home for other defense genes. **A.** A schematic presentation of the *terI* gene locus in *comK*-prophages of various *Lm* strains, showing that the *terI* promoter region is conserved in all prophages. It also shows the different genes that inhabit this locus and their distribution in *comK*-prophages (based on 149 *comK*-prophages; genomes available at NCBI). **B.** A plaque-forming assay of virions obtained from WT *Lm* and *Lm* bacteria expressing *orf80* or *orf180* from the pPL2 plasmid under the regulation of the TetR-dependent promoter (*pPL2-orf80* or *pPL2-orf180*), upon MC treatment (supplemented with 100 ng/ml AT for *pPL2-orf180* culture). Virions were isolated from bacterial lysates 6 h post-MC treatment and tested on the Mack861 indicator strain for PFU. Error bars represent the standard deviation of three independent experiments. **C.** qRT-PCR analysis of the *attB* and *attP* sites in WT *Lm, Lm pPL2-orf80* and *Lm pPL2-orf180* bacteria grown in BHI medium with MC treatment for 3 h, supplemented with AT for *pPL2-orf180* cultures. The results are presented as RQ, relative to the levels in WT bacteria. The experiment was performed three times, and error bars represent the standard deviation of three independent experiments. **D.** qRT-PCR analysis of the *attP* site in encapsidated DNA isolated from virions/capsids collected by ultracentrifugation following DNaseI treatment from WT *Lm* and *Lm pPL2-orf80* or *Lm pPL2-orf180* bacteria (supplemented with AT), upon MC treatment. The experiment was performed three times. Error bars represent the standard deviation of three independent experiments. The upper panel shows Western blot analysis of His_6_-tagged minor capsid protein (LMRG_01534) in virion/capsid samples collected from the corresponding strains (used as a control for the number of virions/capsids). The experiment was performed three times. The figure shows a representative assay. Additional biological repeats can be found in the **Data S1 file**. **E.** A plaque-forming assay of virions obtained upon exogenous infection of *Lm* Δ*ϕ10403S*-Δ*comK,* Δ*ϕ10403S*-Δ*comK+pPL2-orf80* or Δ*ϕ*1*0403S*-Δ*comK+pPL2-orf180* strains with ϕ10403S and ϕEGD857 free phages. Error bars represent the standard deviation of three independent experiments.

## Discussion

This study describes a novel prophage-encoded anti-phage defense system that is prevalent in *L. monocytogenes* prophages, comprising a small defense protein TerI, which inhibits phage DNA packaging, and two self-immunity proteins, anti-TerI1 and anti-TerI2, which counteract TerI activity upon lytic induction of the prophage. The system itself is conserved and positioned within the early lytic gene module, where the self-immunity genes are part of the early lytic operon, and hence transcribed upon lytic induction, whereas the defense gene is constitutively transcribed from the opposite strand (using its own promoter; **Figure 1A**). The findings support a model in which TerI, which is highly produced during lysogeny, targets the terminase complex of invading phages via interaction with TerS, inhibiting interaction of the terminase complex with pre-assembled procapsids, thereby preventing DNA packaging (**Figure 5**). This activity of TerI does not interfere with the other steps of the invading phage lytic cycle, resulting in the release of unpacked, non-infectious procapsids via bacterial lysis driven by the lysis proteins of the invading phage (**Figure 5**). While this mechanism fails to protect infected cells, it confers anti-phage protection to the host (and the prophage) at the population level. The data further indicated that *terI* is also expressed upon lytic induction of the prophage and is fully capable of inhibiting the DNA packaging of its own phage, an observation that led to the discovery of the two self-immunity proteins (anti-TerI1 and anti-TerI2) that counteract TerI activity in the course of ϕ10403S lytic cycle (**Figure 5**). Accordingly, anti-TerI1 and anti-TerI2 were found to be essential for ϕ10403S virion production, as they allowed proper DNA packaging and virion assembly. This study thus uncovered yet another mutualistic interaction between ϕ10403S and its host, describing a unique addition to the growing repertoire of anti-phage defense systems that protect the host without compromising the lytic lifestyle of the phage.

**Figure 5.**
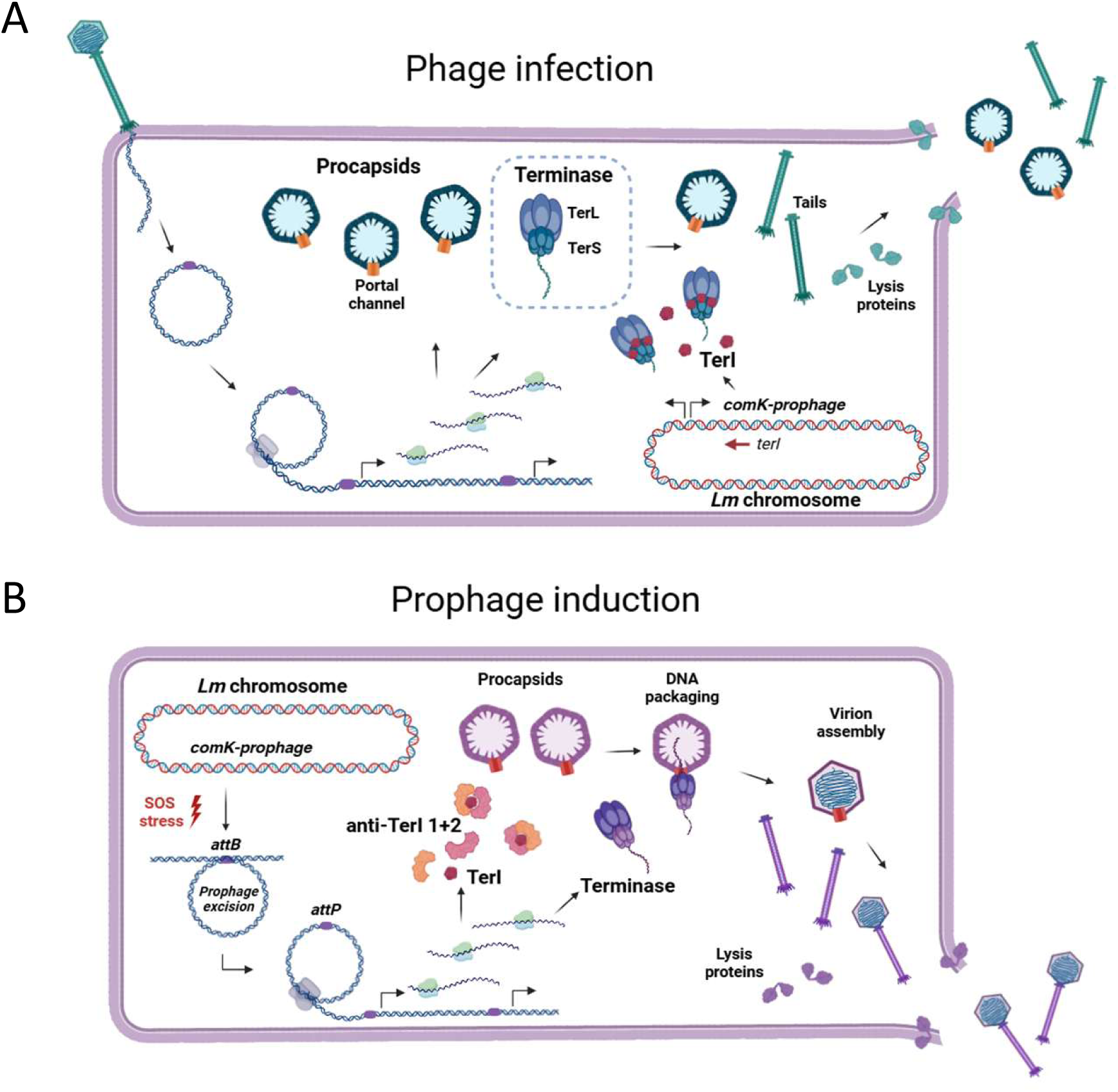
A model of the TERi system: an anti-phage defense system with self-immunity that targets phage DNA packaging. **A.** *terI* is constitutively expressed in lysogenic bacteria from the negative strand of the early genes in the *comK*-prophage. Upon exogenous phage infection, TerI binds the terminase complex of infecting phages via direct interaction with TerS, thereby inhibiting interaction of the terminase with pre-assembled procapsids. This TerI activity prevents phage DNA packaging and virion assembly yet does not interfere with the progression of the infecting phage lytic cycle. Infected cells are eventually lysed by lysis proteins of the invading phage, resulting in the release of non-infectious, unpacked pro-capsids that cannot infect neighboring cells. While this system does not protect infected cells, it provides anti-phage protection to the host (and the prophage) at the level of the population. **B.** Upon induction of the *comK*-prophage, both TerI and the self-immunity proteins anti-TerI1 and anti-TerI2 are expressed. anti-TerI1 and anti-TerI2, which are encoded by genes belonging to the phage early lytic gene module, counteract TerI activity via direct interaction, allowing the *comK-*phage to package its DNA and assemble progeny virions, which are then released into the environment via bacterial lysis.

Phage DNA packaging is a highly coordinated process mediated by several proteins. In tailed phages, it is thought to be facilitated by a molecular motor composed of three proteins, the portal protein that forms the channel through which DNA enters the procapsid, the small terminase subunit (TerS) that recognizes phage DNA and the large terminase subunit (TerL), which translocates such DNA into the procapsids using energy derived from ATP hydrolysis (using the TerL ATPase domain).^30–34^ Upon completion of DNA packaging, TerL further cuts the DNA and dissociates from the nascent virion, leaving behind the portal channel to recruit other proteins that stabilize the packaged DNA and plug the channel, with the portal remaining an integral part of the virion.^34^ As noted, during the process of DNA translocation, the procapsid shell expands to accommodate the complete phage genome and is then decorated by additional proteins.^31,35^ Here, we found that TerI directly interacts with TerS, which is the component that initiates the DNA packaging process. It is thought that TerS first binds the phage DNA and then recruits TerL, triggering the formation of TerS-TerL complexes that subsequently bind pre-assembled procapsids via the portal, with such interactions being mediated by TerL.^30,36^ As such, terminase complexes can be found free in infected cells or bound to procapsids while involved in DNA translocation.^37,38^ Co-sedimentation experiments indicated that TerI is not associated with procapsids, excluding the possibility that TerI targets the portal channel of nascent virions or directly interferes with DNA translocation. Instead, the data demonstrated that TerI targets the terminase complex, likely in its free form, i.e., before it associates with a nascent virion, thus inhibiting its interaction with the portal to prevent DNA packaging. TerI activity led to the accumulation of small procapsids of 40 nm devoid of DNA, resembling procapsids observed in bacteria that lack the terminase complex (i.e., D*terSL* bacteria). These findings support the premise that TerI is a direct inhibitor of the terminase complex, having evolved to inhibit DNA packaging as an anti-phage defense. That said, it remains unclear how TerI activity is directly manifested. Does it affect the assembly or the multimeric composition of the terminase complex (which comprises two multimeric rings, one of TerS and the other of TerL) or does it directly prevent terminase complex interaction with the procapsid? Questions that require further investigation.

The field of anti-phage defense has seen remarkable advances in recent years, with the discovery of multiple systems that provide anti-phage immunity. Bacteria were shown to encode a variety of systems that protect from phage predation, such as restriction-modification (RM), Abi and adaptive immune systems, such as CRISPR-Cas.^39–44^ In addition, prophages and their derivatives, including cryptic prophages, phage satellites and other mobile genetic elements, which inhabit bacterial genomes, were also shown to carry defense genes, implying that phage resistance is a common interest.^7–9,45,46^ It is believed that phage-encoded systems are used to protect prophages from inter-phage competition and predation, thus playing an important role in the evolution of temperate phages and their hosts. Phage-encoded systems were shown to comprise various mechanisms, such as Sie systems that directly prevent phage infection by inhibiting phage entry, thereby providing cell protection, RM and CRISPR-Cas systems that target incoming phage DNA upon injection, thus allowing the infected cells to survive, and Abi systems that induce cell death or growth arrest, sacrificing infected cells for the benefit of the rest by preventing epidemic spread of the phage within the population.^7–9,45,46^ Another mechanism that emerged from recent reports resembles that described here for TERi. This mechanism involves direct inhibition of virion assembly, while allowing bacterial lysis by the infecting phage to proceed. Two independent studies have demonstrated phage-encoded anti-phage defense systems that specifically inhibit tail assembly without preventing bacterial lysis. The Tab (tail assembly blocker) system of *Pseudomonas aeruginosa* prophages^47^ and the Tai (tail assembly inhibitor) system of *Escherichia coli* and *Salmonella enterica* prophages^48^ were shown to target tail tape-measure protein and central tail fibres, respectively, preventing tail assembly and hence virion production. Of note, these systems were also demonstrated to include counter-defense proteins that allow proper virion assembly of the cognate phage. The Tab and Tai systems share common features with TERi as they all target virion assembly while allowing phage-mediated bacterial lysis by the invading phage, leading to the release of tailless phage particles or empty procapsids that cannot infect neighbouring cells. As indicated, the existence of dedicated self-immunity determinants is also shared, demonstrating the requirement for a mechanism that prevents auto-immunity, given how the defense proteins target conserved processes of phage assembly. Moreover, the genetic architecture of the TERi system mirrors that of the Tab and Tai systems in that defense and counter-defense genes are transcribed from opposite strands. This allows the counter-defense proteins to be produced upon prophage induction and inactivate the defense protein in the course of the lytic cycle. This genetic arrangement exemplifies how prophages managed to acquire such defense systems into their gnomes, by aligning their expression with the different lifestyles of the phage.

Similar defense/counter-defense mechanisms have been demonstrated in other prophage-encoded systems, such as the Tha system of *Staphylococcus* prophages and the BstA system of *Salmonella enterica* prophage.^49,50^ In these systems, the Tha and the BstA defense proteins trigger abortive infection upon recognition of a tail protein or phage DNA replication, respectively. Moreover, these systems further prevent auto-immunity by expressing counter-defense determinants during prophage induction. In the case of the Tha system, the counter-defense protein Ith was shown to be encoded by a gene located nearby, yet on the opposite strand, creating a non-contiguous operon with its cognate defense gene.^49^ In the BstA system the counter-defense determinant was shown to be a DNA element (aba), whose mode of action remains unclear.^50^ Indeed, in all of these cases, the defense systems could not exist in the absence of the counter-defense genes.

Inhibition of virion assembly as a defense strategy is not unique to prophages. A recent report demonstrated that *Collimonas* sp. OK412 bacteria encode the Bil system that inhibits tail assembly of invading phages.^51^ This system was shown to conjugate bacterial ubiquitin-like proteins to the phage central tail fibres, leading to the release of tailless phage particles that cannot infect neighboring cells. Another example is the BREX system identified in *Bacillus cereus*, which inhibits phage DNA replication via a mechanism that is still unclear but involves DNA methylation.^52^ The AbiZ system, which is encoded by a conjugative plasmid of *Lactococcus* spp. was shown to indirectly inhibit virion assembly by triggering early bacterial lysis of infected cells (possibly via interaction with the phage holin), thereby leading to the release of immature phage particles.^53^ In this regard, the Tab, Tai, Bil and TERi systems also result in the release of non-assembled phage particles, with cell lysis being mediated by the lysis proteins of the invading phage. Of note, phage-mediated bacterial lysis is a process that is regulated independently of virion assembly, as it occurs regardless of whether progeny virions are successfully assembled.^54^ These systems can thus be considered as guaranteeing that no progeny virions form and that infected cells die, ultimately blocking phage epidemics from spreading within the population. While this outcome is similar to that elicited by Abi systems, the main difference between the two strategies is that Abi systems rely on proteins that sense phage infection and trigger growth arrest or cell death, thus indirectly preventing virion production.^10^ In contrast, the TERi, Tab and Tai systems rely on proteins that directly interfere with virion assembly while allowing bacterial lysis by the infecting phage. Both strategies provide anti-phage defense at the population-level, a tactic that appears to be successful, as evidenced by the various bacteria and phage-encoded such defense systems that have been discovered so far.

The targeting of the terminase complex by TerI reveals an interesting parallel with phage satellite elements, like phage-inducible chromosomal islands (PICIs) of *Staphylococcus aureus*, which have long been known to manipulate the packaging machinery of helper phages to promote their dissemination.^55^ PICIs were shown to encode TerS homologs and Ppi proteins that specifically block TerS of the helper phage to facilitate PICI packaging instead of phage DNA.^56,57^ Another such example is the PICI-encoded protein RppA, which was shown to direct the TerS of a helper phage to recognize the PICI genome and facilitate its packaging.^58^ The phage satellite of *Vibrio cholera* PLE was also shown to target the terminase complex of another phage. PLE encodes the Gpi protein that targets TerL of the lytic phage ICP1, thereby hijacking the phage packaging machinery to promote its horizontal spread.^59^ While Ppi, RppA and Gpi evolved to facilitate the dissemination of their elements, TerI interaction with the terminase complex represents a distinct evolutionary solution that has evolved for defense. In this study, we identified two additional proteins that target phage DNA packaging, namely TerI-80 and TerI-180, indicating that inhibition of DNA packaging as a defense is widespread.

The ability of temperate phages to compete with other phages and protect their bacterial hosts from predation represents a sophisticated form of symbiosis. This relationship contributes to the stability of lysogenic interactions and explains the widespread prevalence of prophages in bacterial genomes. Understanding bacteria-phage interactions and the role of anti-phage defense systems in shaping bacterial communities and the evolution of pathogenic strains may lead to the development of new therapeutic approaches to treat bacterial infections (e.g., using phage therapy) and control pathogen propagation in natural environments.

## Materials and methods

### Bacterial strains, phages, plasmids and growth conditions

*L. monocytogenes* strain 10403S was used as WT strain and the parental strain for generating all mutants in this study unless indicated otherwise. *E. coli* XL-1 Blue (Stratagene) was utilized for vector propagation. *E. coli* SM-10 was utilized for conjugative plasmid delivery to *L. monocytogenes* bacteria. *Listeria innocua* CLIP11262 was used to amplify *orf80,* and A006 phage was used to amplify *orf180*. *Listeria* strains were grown in brain heart infusion (BHI) (Merck) medium at 37°C or 30°C as specified, and *E. coli* strains were grown in Luria-Bertani (LB) (Acumedia) medium at 37°C. *Listeria* phages ϕEGD857, ϕC118, ϕA006, ϕA511 and ϕP100 used in this study were a gift from Richard Calendar (UC Berkeley) and Joe Bondy-Denomy (UCSF). The phages were propagated in *L. monocytogenes* strain Mack861 and kept in 10% DMSO at - 80°C. Phusion DNA polymerase was used for all cloning purposes, and *Taq* polymerase was used for PCR to verify the different plasmids and strains. Antibiotics were used as follows: chloramphenicol (Cm), 10 µg/ml; streptomycin (Strep), 100 µg/ml; kanamycin (Km), 30 µg/ml; and mitomycin C (MC) (Sigma), 1.5 µg/ml. All restriction enzymes were purchased from New England BioLabs.

### Generation of gene deletion mutant and over-expression strains

To generate gene deletion mutants, upstream and downstream regions of the selected gene were amplified using Phusion DNA polymerase and cloned into the pBHE plasmid (pKSV-oriT). The plasmids were then verified by PCR and their inserts were sequenced. Plasmids were then conjugated to *L. monocytogenes* using *E. coli* SM-10 bacteria. *Trans-*conjugants were selected on BHI agar plates supplemented with chloramphenicol and streptomycin and transferred to BHI medium supplemented with chloramphenicol for two days at 41°C to allow plasmid integration into the bacterial chromosome by homologous recombination. The bacteria were passed several times in fresh BHI medium without chloramphenicol at 30°C to promote plasmid loss (curing) and the generation of an in-frame gene deletion. The bacteria were plated on BHI plates with or without chloramphenicol, and sensitive colonies were validated for gene deletion by PCR. For over-expressing strains, the corresponding gene was cloned into the pPL2 integrative plasmid under regulation of the TetR-dependent promoter.^11^ The PCR-amplified insert was sequenced for verification, and the cloned plasmid was conjugated to *L. monocytogenes* using *E. coli* SM-10 bacteria. The TetR inducer anhydrotetracycline (AT) was supplemented at a concentration of 100 ng/ml to induce enhanced gene expression where indicated.

### Bacterial growth and prophage induction

*Lm* bacteria were grown overnight (O.N.) in BHI medium at 37°C with agitation. To examine lysogenic bacteria, an O.N. culture was diluted by a factor of 100 in fresh BHI and incubated without agitation at 30°C to reach an OD_600nm_ of 0.7-0.8. For induction of the prophage, UV irradiation or MC treatment was performed. For UV irradiation, O.N. cultures were diluted by a factor of 10 in 10 ml of fresh BHI medium and incubated without agitation at 30°C to reach an OD_600nm_ of 0.5. The cultures were irradiated by UV light at 4 J/cm² (using a CL 508S model UV cross-linker oven), supplemented with 5 ml of fresh BHI medium, and incubated without agitation at 30°C. For induction of the prophage by MC, bacteria were grown O.N. at 37°C with agitation in BHI medium, diluted by a factor of 10 ml in fresh BHI, incubated without agitation at 30°C to reach an OD_600nm_ of 0.4-0.5, diluted again to an OD_600nm_ of 0.15 and supplemented with MC (1.5 µg/ml) for 6 h. To follow bacterial growth or lysis (e.g., under lytic conditions), bacteria were grown in 96-well plates, incubated at 30°C in a Synergy HT BioTek plate reader and monitored for OD_600nm_ every 15 min after 2 min of shaking. Experiments were repeated at least three times.

### Plaque-forming assay to assess phage induction and infection

To analyze virion production upon prophage induction, bacteria were grown O.N. at 37°C with agitation in BHI medium, diluted by a factor of 10 in fresh BHI and incubated without agitation at 30°C to reach OD_600nm_ of 0.4. The cultures were then diluted again to an OD_600nm_ of 0.15, supplemented with MC (1.5 µg/ml), and incubated at 30°C for 6 hours without agitation. The treated cultures were then filtered through 0.22 μm filters to allow passage of virions but not bacteria. Serial dilutions of filtered supernatants (100 μl) were added to 3 ml of melted 0.7% agar-containing LB medium (56°C), and 300 μl of an O.N. culture of *L. monocytogenes* strain Mack861, which served as an indicator strain, and quickly overlaid on BHI-agar plates. The plates were then incubated at room temperature for 3 to 4 days to allow plaque formation. To analyze virion production upon exogenous phage infection, free phage particles (100 μl) were used to infect *Δϕ10403S/ΔcomK* bacteria in 3 ml of melted 0.7% agar-containing LB medium (56°C), which was then quickly overlaid on BHI-agar plates and analyzed as described above.

### qRT-PCR and RT-qPCR analyses

Bacteria were grown at 37°C O.N. in BHI broth with agitation, diluted 1:10 in BHI, incubated without agitation at 30°C to reach an OD600nm of 0.4, diluted to an OD600nm of 0.15, at which point a lytic cycle was induced by addition of MC (1.5 µg/ml). Bacteria were harvested by centrifugation at indicated time points and snap-frozen in liquid nitrogen. Total nucleic acids were isolated using standard phenol-chloroform extraction. For analysis of *attB/attP* levels by qRT-PCR, 0.04 ng total nucleic acids were used, with the bacterial 16S rRNA gene being used as a reference for sample normalization. For gene transcription analysis (RT-qPCR), samples were treated with DnaseI, and 1 μg RNA was reverse-transcribed to cDNA using a qScript (Quanta) kit. For strand specific analysis, 1 μg of RNA was reverse transcribed to cDNA using a reverse primer of the particular gene of interest and a qScriptFlex (Quanta) kit. RT-qPCR was performed on 10 ng cDNA. The relative expression of bacterial genes was determined by comparing their transcript levels with those of the bacterial 16S rRNA gene, which served as reference. All RT-qPCR analyses were performed using PerfeCTa SYBR Green FastMix (Quanta) on the StepOnePlus RT-PCR system (Applied Biosystems), per the standard ^ΔΔ^C_t_ method. Experiments were performed in three biological replicates.

### Transmission electron microscopy

To visualize virion production using TEM, bacteria were harvested at 4 hours post-UV irradiation and fixed with 2.5% glutaraldehyde in PBS overnight at 4°C. After several washes with PBS, the cells were treated with 1% OsO_4_ (in PBS) for 2 hours at 4°C. Dehydration was carried using a graded ethanol series followed by embedding in glycidyl ether. Thin sections were mounted on Formvar/Carbon coated grids, stained with uranyl acetate and lead citrate and examined using a Jeol 1400 – Plus transmission electron microscope (Jeol, Japan). Images were captured using SIS Megaview III and iTEM, the Tem imaging platform (Olympus).

### Analysis of encapsidated phage DNA

MC-treated bacterial cultures were filtered through a 0.22 μm filter, and virions/capsids from 1 ml of the filtrate were collected by centrifugation at 100,000 g for 10 minutes using an Optima max XP centrifuge. The supernatant was then removed, and the pellet was suspended in 50 μl of DNaseI buffer, supplemented with 1 μl of DNaseI (Thermo Scientific), and incubated at 37°C for 1 h to degrade external DNA. Denaturation of virions/capsids was performed using a standard solution of phenol-chloroform, and the released phage DNA (encapsidated DNA) was supplemented with the same amount of a plasmid encoding the *Km* resistance gene, which was used to normalize the samples for qRT-PCR. DNA samples were extracted using a standard phenol-chloroform-based procedure followed by ethanol precipitation. Purified DNA was resuspended in 100 μl of water and directly analyzed by qRT-PCR to determine the amounts of the *attP* site and the *Km* gene.

### Western blot analysis of ORF65

Indicated strains were grown in 50 ml BHI at 37°C to OD_600nm_ of 0.6. The cultures were then harvested and washed with buffer A (20 mM Tris-HCl, pH = 8, 0.5 M NaCl, and 1 mM EDTA), resuspended in 1 ml of buffer A supplemented with 1 mM phenylmethylsulfonyl fluoride (PMSF) and lysed by ultra-sonication. Total protein content was assayed using a modified Lowry assay and aliquots containing equal amounts of total protein were separated on 15% SDS-polyacrylamide gels and transferred to nitrocellulose membranes. Proteins were probed with rabbit anti-His_6_ tag (Abcam ab9108) antibodies used at a 1:1000 dilution, followed by HRP-conjugated goat anti-rabbit IgGs (Jackson ImmunoResearch) at a 1:20,000 dilution. Western blots were developed using a homemade enhanced chemiluminescence (ECL) reagent.

### Analysis of ORF65 sedimentation with phage particles

For analysis of ORF65 association with phage particles/complexes, bacteria were grown O.N. in BHI medium at 37°C with agitation and then diluted by a factor of 10 ml in fresh BHI and incubated without agitation at 30°C to reach an OD_600nm_ of 0.4. The cultures were again diluted to an OD_600nm_ of 0.15 and prophage induction was triggered using MC (1.5 µg/ml) for 6 hours. Bacterial lysetes were then filtered through 0.22 μm filters that allow passage of phage particles but not bacteria. The filtrate (25 ml) was centrifuged at 100,000 g for 30 minutes using a Beckman Coulter Optima L-90K centrifuge. The supernatant was removed, and the pellet was suspended in 100 μl of buffer A. To detect His_6_-tagged ORF65 in supernatant fractions, the proteins were concentrated using standard trichloroacetic acid (TCA) precipitation and suspended in 100 μl of 1% SDS. The pellet or supernatant samples were then separated on 15% SDS-polyacrylamide gels and transferred to nitrocellulose membranes for Western blot analysis using rabbit anti-His_6_ tag antibodies (Abcam ab9108-1:1000 dilution), followed by HRP-conjugated goat anti-rabbit IgGs (Jackson ImmunoResearch; 1:20,000 dilution). The amount of (pro)capsids was detected using anti-His_6_ tag antibodies against a His_6_-tagged minor capsid protein (LMRG_01534). Western blots were developed using a homemade ECL reagent.

### Bacterial adenylate cyclase-based two-hybrid system

The assay was performed using a Bacterial Adenylate Cyclase Two-Hybrid (BACTH) system (Euromedex), according to the manufacturer’s instructions. The *orf65* gene was cloned into the pUT18C plasmid, and the *LMRG_02984*, *LMRG_01518*, *terS* and *terL* genes were cloned into the pKT25 plasmid. *E. coli* BTH101 electrocompetent cells were co-transformed with the two plasmids, plated on LB plates supplemented with ampicillin (100 µg/mL), kanamycin (30 µg/mL) and IPTG (0.5 mM), and incubated for 48 hours at 30°C. To detect protein-protein interactions, overnight LB cultures were grown at 37°C in the presence of ampicillin (100 µg/mL), kanamycin (30 µg/mL) and IPTG (0.5 mM), and the assay for β-galactosidase activity was performed using *ortho*-nitrophenyl-β-galactoside (ONPG, Sigma-Aldrich), according to the manufacturer’s instructions. The enzymatic reaction was followed using Synergy HT Biotech plate reader for 30 min at room temperature, measuring 420 nm absorbance every 2 min. β -galactosidase activity was calculated based on two proximal time points in the linear part of the curve, and normalized to the OD_600nm_ of the sample. The experiments were performed in triplicate and independently repeated three times.

### Suppressor mutation screen

To identify suppressor mutations that restore virion production in the presence of TerI, cultures of WT bacteria harboring the ϕ10403S prophage with a kanamycin resistance gene^11^ with or without plasmid pPL2*-terI* were subjected to UV irradiation. Bacterial lysates were filtered through a 0.2 μm filter and assayed for plaque formation. Fifty plaques appeared from bacteria expressing *terI*, as opposed to 10^6^ PFUs that appeared in the control strain. Virions from the 50 plaques were picked and incubated O.N. in 1 ml of PBS at 4°C. Each sample was treated with chloroform and then filtered through a 0.22 μm filter. Next, fresh *Δϕ10403S/ΔcomK* bacteria expressing *terI* were infected with the virion filtrate of each potentially mutated phage at MOI of ∼1:100 and incubated O.N. at 30°C, allowing the phages to lysogenize into the bacterial chromosome at *comK*. Lysogens were selected on BHI plates supplemented with Km. Fresh cultures of lysogenized bacteria were then UV irradiated, and virions were isolated again from the plaques, and used to infect fresh *Δϕ10403S/ΔcomK* bacteria expressing *terI*. At each round of phage infection, titers of infective virions produced by phage mutants were monitored and compared to the WT phage. This experiment was performed in four separate lines, with each line undergoing 10 rounds of phage induction and lysogenization. Seventeen phage mutants were isolated from the same line. Lysogens carrying the suppressor phage mutants were sequenced at the Technion Genome Centre (Haifa, Israel).

### Construction of pSNIB1 integrative plasmid

The pSNIB1 integrative plasmid was generated on the backbone of plasmid pPL2 using the integrase and the attachment site of ϕLMC1 prophage, which integrates into the tRNA^Ser^ gene. The ϕLMC1 integrase and attachment site were synthesized and cloned into a BlueScript vector by GeneScript, and amplified by PCR. The pPL2 plasmid was digested with SphI and BglII restriction enzymes, separated on agarose gel, purified and used as a vector for Gibson assembly with the PCR product. Cloned plasmids were verified by PCR and their PCR-amplified inserts were sequenced. The plasmids were conjugated to *L. monocytogenes* using *E. coli* SM-10 bacteria. *Trans*-conjugants (WT-pSNIB1-Km) were selected on BHI agar plates supplemented with chloramphenicol and streptomycin. For analysis of plasmid integration, PCR was performed on three *trans*-conjugant colonies, amplifying *attL* and *attR* sites. The chloramphenicol resistance gene was then replaced with the kanamycin resistance gene using PvuI and ApaLI restriction enzymes.

### Electromobility shift assay

To purify Cro, the *cro*-like gene (*LMRG_01515*) was amplified from WT *Lm* by PCR using primers containing DNA encoding a His_6_-tag directed to the C-terminus. The purified PCR fragment was cloned into plasmid pET11a using NdeI and BamHI restriction enzymes, and the PCR-amplified insert was verified by sequencing (pET11a-*cro-6his*). *E. coli* BL21(λDE3) expressing Cro-His_6_ were grown in 500 ml of LB medium supplemented with 50 µg/ml ampicillin at 37°C to an OD_600nm_ of ∼0.6. The cultures were then induced with 0.5 mM IPTG, and grown for 3 hours at 32°C. Cells were harvested by centrifugation and washed with 0.9% NaCl. Cro-His_6_ was purified as described^60^ with some modifications. Cells were lyzed by sonication (70% amplitude, 9 s pulses) in 10 ml of lysis buffer (50 mM sodium phosphate (pH 7.6), 500 mM NaCl, 5% glycerol, 10 mM imidazole, 1 mM PMSF) and cell debris were removed by centrifugation at 16,000 g for 45 min at 4°C. The supernatant fractions were loaded onto 2 ml column of Ni-NTA resin (Thermo Fisher Scientific) saturated with the lysis buffer and washed thoroughly with 20 bead volumes of wash buffer (50 mM sodium phosphate (pH 7.6), 500 mM NaCl, 5% glycerol, 25 mM imidazole). Protein fractions were eluted using elution buffer (50 mM sodium phosphate (pH 7.6), 500 mM NaCl, 5% glycerol, 250 mM imidazole) and analyzed by 12% SDS–PAGE. Fractions containing the His-tagged protein were dialyzed in a stepwise manner using buffer D1 (50 mM sodium phosphate (pH 7.6), 400 mM NaCl, 5% glycerol, 1 mM EDTA), followed by buffer D2 (50 mM sodium phosphate (pH 7.6), 200 mM NaCl, 5% glycerol, 1 mM EDTA), and finally buffer D3 (50 mM sodium phosphate (pH 7.6), 400 mM NaCl, 5% glycerol). Protein concentration was estimated using a standard Bradford method (Bio-Rad).

For gel shift assays, a duplex DNA sequence carrying 16 bp of the putative *Cro* operator site with six flanking nucleotides at each end (TTATAG<u>AGAAACAAAACGTTTC</u>TTTTGG), as well as a mutated operator DNA duplex (TTATAG<u>AGAAACAAAACGTTTC</u>TTTTGG) were purchased from Integrated DNA Technologies. Increasing concentrations of Cro protein (0-1 µM) were incubated with 10 nM of probe DNA in buffer D3 for 20 min on ice, as described previously.^60,61^ The samples were then analyzed using 10% native PAGE. The gels were stained with SYBR Green II (Thermo Fisher Scientific) and visualized using an Odyssey M Imager. To analyze Cro binding to its putative operator site, ImageJ was used with the “plot profile” function.^62^ The experiment was performed three times.

### Genome analysis of *Listeria comK*-prophages

A number of complete *Listeria* genus and individual *Listeria* species genomes were retrieved from the NCBI Genomes database (https://www.ncbi.nlm.nih.gov/genome, accessed on November 1, 2024). Particularly, the Genome Assembly and Annotation report (https://www.ncbi.nlm.nih.gov/genome/browse/#!/prokaryotes/159/; accessed on January 1, 2024) website provides a list of both complete genomes and incomplete genome assemblies of *Listeria monocytogenes* strains. The above-mentioned website also provides a filter option by which one can exclusively analyze complete *L. monocytogenes* genomes (“Chromosome” and “Complete” should be selected). This approach revealed a number of complete genomes of other *Listeria* species (e.g. *L. innocua*, *L. seeligeri*, *L. ivanovii*). To determine the number of complete *Listeria* genomes that carry intact *comK*-associated prophages, we developed an approach based on a *comK*-phage-specific query (the integrase gene sequence of the *Listeria* phage A118) and the customized Nucleotide BLAST machine setup at the NCBI site (https://blast.ncbi.nlm.nih.gov/Blast.cgi; accessed on November 1, 2024), using the following filters: i) Nucleotide collection (nr/nt) database (consists of GenBank+EMBL+DDBJ sequences, excluding WGS data); ii) Max target sequences (500); and iii) Expect threshold (either 10 or 100). This search yielded a list of complete genomes of *Listeria* species, as well as the two known *Listeria* phages, A118 and PSU-VKH-LP019, that possess an entire sequence of the integrase gene. A similar approach was used to verify a number of complete *Listeria* genomes using the nucleotide sequence of either the *dnaA* or *dnaN* gene of the reference *L. monocytogenes* strain 10403S. As of January 2024, 422 complete *Listeria* genomes have been identified, of which 149 were found to carry a prophage in the *comK* gene. From these genomes, the sequences of *comK*-prophages were extracted and further analyzed using standard tools available online.

## Acknowledgements

This work was supported by the European Research Council (ERC) consolidator grant (Co-Patho-Phage 817842) and the Israel Science Foundation (ISF 1599/23) for A.A. Herskovits. We thank Joe Bondy-Denomy from UCSF for sending us *Listeria* phages.

## Author contributions

G. Azulay, N. Sigal and A.A. Herskovits designed the study. G. Azulay, N. Sigal, K. Tzohar Rabinovich, O. Stadnyuk, A. Das, P. Lisitzin and E. Biderman performed the experiments. N. Sigal validated the results. D. Koenigsfeld and I. Borovok performed bioinformatic analysis, and A.A. Herskovits wrote the manuscript.

---

**Figure S1.**
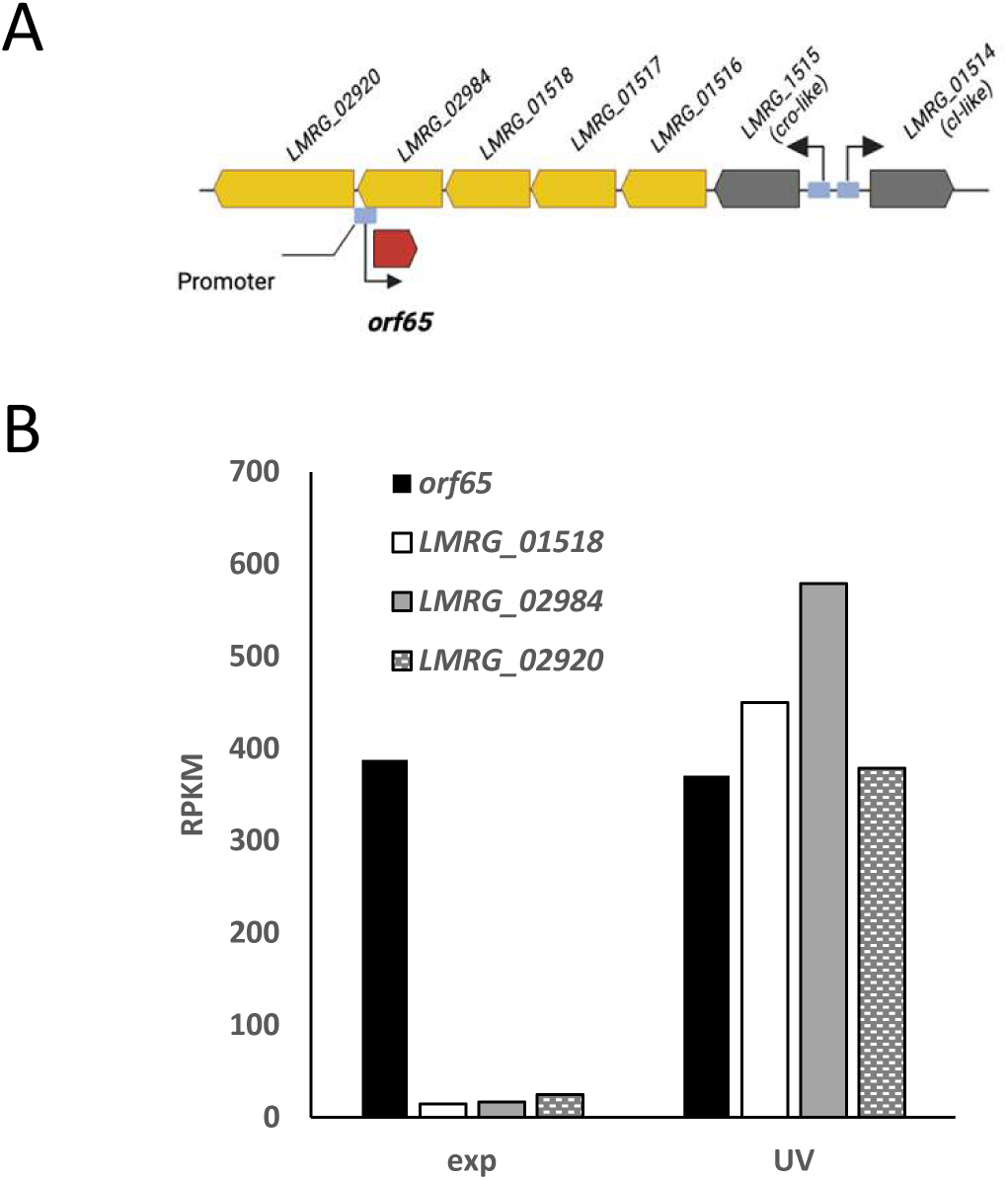
*orf65* is expressed during lysogeny and upon prophage induction. **A.** Schematic representation of the *orf65* genomic locus. **B.** Transcription analysis of the *orf65* genomic locus in *Lm* 10403S bacteria irradiated with UV light (4 h post-UV irradiation) or not, as detected by strand-specific RNA-seq analysis. Bacteria were grown in BHI medium to exponential phase at 30°C. Transcription levels are indicated as RPKM (scaled reads per kilobase per million reads). Data represent the mean of three independent experiments.

**Figure S2.**
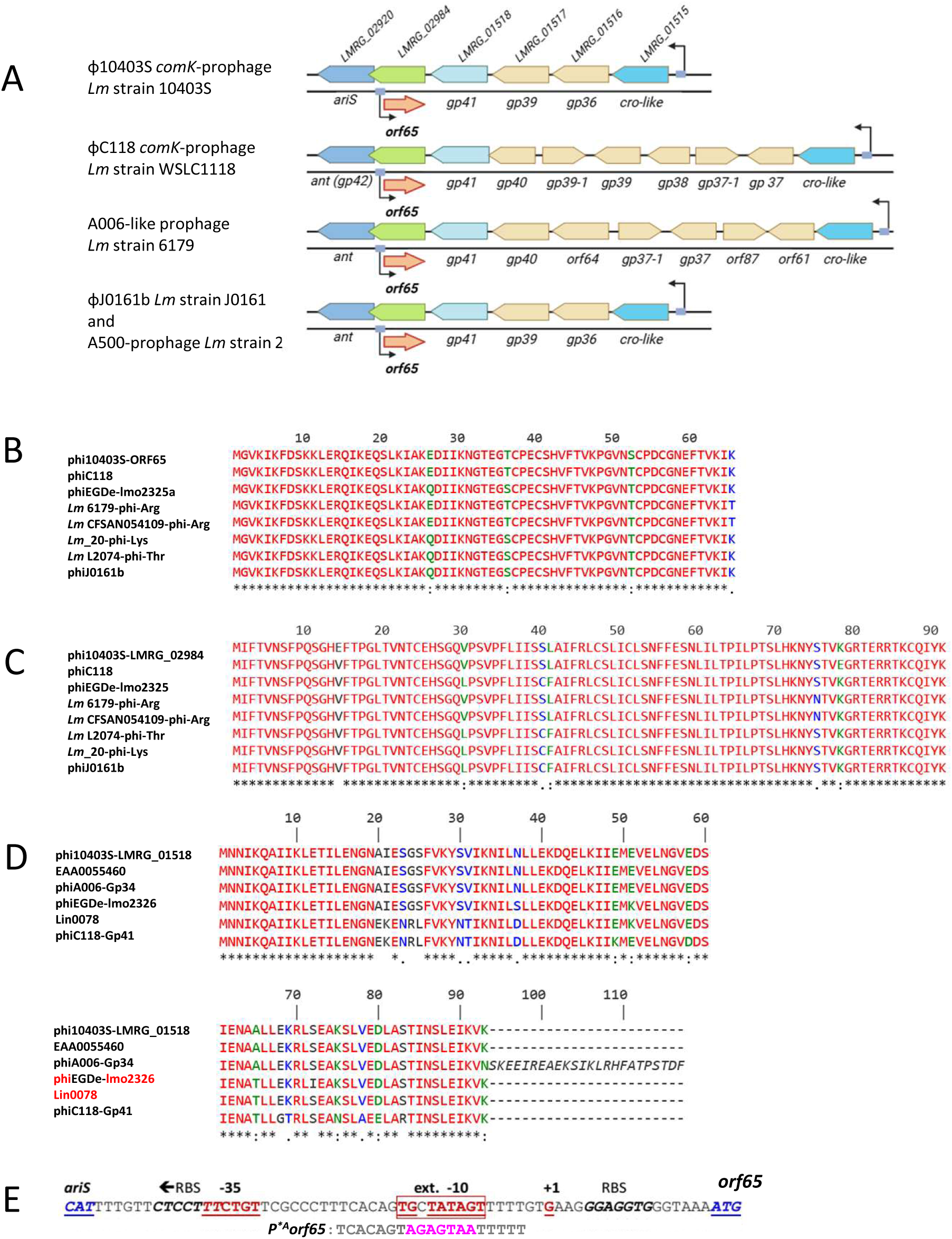
ORF65 is conserved in *Listeria* phages. **A.** Schematic representation of the genomic region of ORF65 homologues in different *Listeria* prophages. **B.** Conservation of ORF65. Multiple alignment of ORF65 and its homologues. phi10403S-ORF65 of the *comK-*prophage of *Lm* strain 10403S (not annotated in CP002002.1); phiC118, ORF65 ortholog of *comK*-prophage of *Lm* strain WSLC1118; phiEGDe-lmo2325a, ORF65 ortholog of *comK-*prophage of *Lm* strain EGD-e (locus tag lmo2325a in NC_003210.1); *Lm* 6179-phi-Arg, ORF65 ortholog of the A006-like prophage of *Lm* strain 6179 (locus tag LM6179_1752, accession CDM20015.1); *Lm* CFSAN054109-phi-Arg, ORF65 ortholog of the A006-like prophage of *Lm* strain CFSAN054109 (locus tag C9J84_13600); *Lm*_20-phi-Lys, ORF65 ortholog of the A500-like prophage of *Lm* strain 20 (not annotated in CP030803.1)*; Lm* L2074-phi-Thr, ORF65 ortholog of the tRNA-Thr associated prophage of *Lm* strain L2074 (locus tag L2074_02706); phiJ0161b, ORF65 ortholog of the tRNA-Thr associated prophage of *Lm* strain J0161 (not annotated in CP002001.1). **C.** Conservation of LMRG_02984. Multiple alignments of LMRG_2984 and its homologues. phi10403S-LMRG_02984 of the *comK-* prophage of *Lm* strain 10403S; phiC118, LMRG_02984 ortholog of the *comK-pro*phage of *Lm* strain WSLC1118; phiEGDe-lmo2325, LMRG_02984 ortholog of the *comK-*prophage of *Lm* strain EGD-e (locus tag lmo2325 in NC_003210.1); *Lm* 6179-phi-Arg, LMRG_02984 ortholog of the A006-like prophage of *Lm* strain 6179 (not annotated in HG813249.1); *Lm* CFSAN054109-phi-Arg, LMRG_02984 ortholog of the A006-like prophage of *Lm* strain CFSAN054109 (not annotated in CP028183.1); *Lm* L2074-phi-Thr, LMRG_02984 ortholog of the tRNA-Thr associated prophage of *Lm* strain L2074 (not annotated in CP007689.1); *Lm*_20-phi-Lys, LMRG_02984 ortholog of the A500-like prophage of *Lm* strain 20 (locus tag DRA62_14180); phiJ0161b, LMRG_02984 ortholog of the tRNA-Thr associated prophage of *Lm* strain J0161 (locus tag LMOG_02942). **D.** Conservation of LMRG_01518. Multiple alignment of LMRG_1518 and its homologues. phi10403S-LMRG_01518 of the *comK-*prophage of *Lm strain* 10403S; EAA0055460 of the tRNA-Thr associated prophage of *Lm* FDA1045857-026-002 (accession number EAA0055460.1; locus tag EEP75_15435); phiA006-Gp34, LMRG_02984 ortholog (Gp34) of the *Listeria* phage A006 (harboring *orf180*, annotated as *gp35*, in place of *orf65*); phiEGDe-lmo2326, LMRG_02984 ortholog of the *comK-*prophage of *Lm* EGD-e (locus tag lmo2326 in NC_003210.1), Lin0078, LMRG_02984 ortholog of the A500-like prophage of *L. innocua* Clip 1126293 (harboring *orf80*, annotated as *lin0079*, in place of *orf65*); phiC118-Gp41, LMRG_02984 ortholog of the *comK-*prophage of *Lm* strain WSLC1118. Amino acid sequences were aligned using CLUSTALW (https://npsa-prabi.ibcp.fr/cgi-bin/npsa_automat.pl?page=/NPSA/npsa_clustalw.html); identical residues marked with asterisks, highly conserved residues with colons, and weakly conserved residues with dots. **E.** The promoter region of the *orf65* gene. The P^*A^*orf65* mutation is marked in magenta.

**Figure S3.**
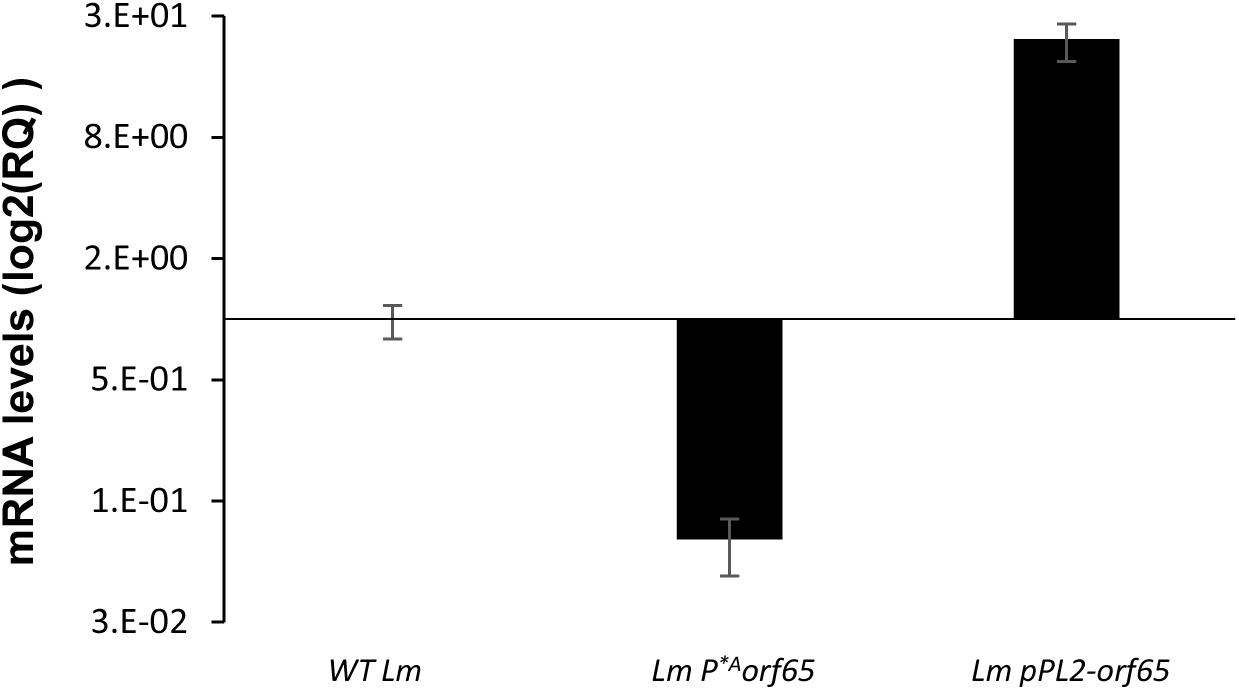
*orf65* transcription levels in different strains. RT-qPCR analysis of *orf65* transcription levels in WT *Lm, Lm* P^*A^*orf65* (harboring a mutation in *orf65* promoter) and *Lm* bacteria ectopically expressing *orf65* from the integrative pPL2 plasmid under regulation of the constitutive *rpsD* promoter (pPL2-*orf65*). The strains were grown exponentially in BHI medium at 30°C. mRNA levels are presented as a log of their relative quantity (RQ), relative to their levels in WT bacteria. The experiment was performed three times. The figure shows a representative assay, with error bars representing a 95% confidence interval. Additional biological repeats can be found in the **Data S1 file**.

**Figure S4.**
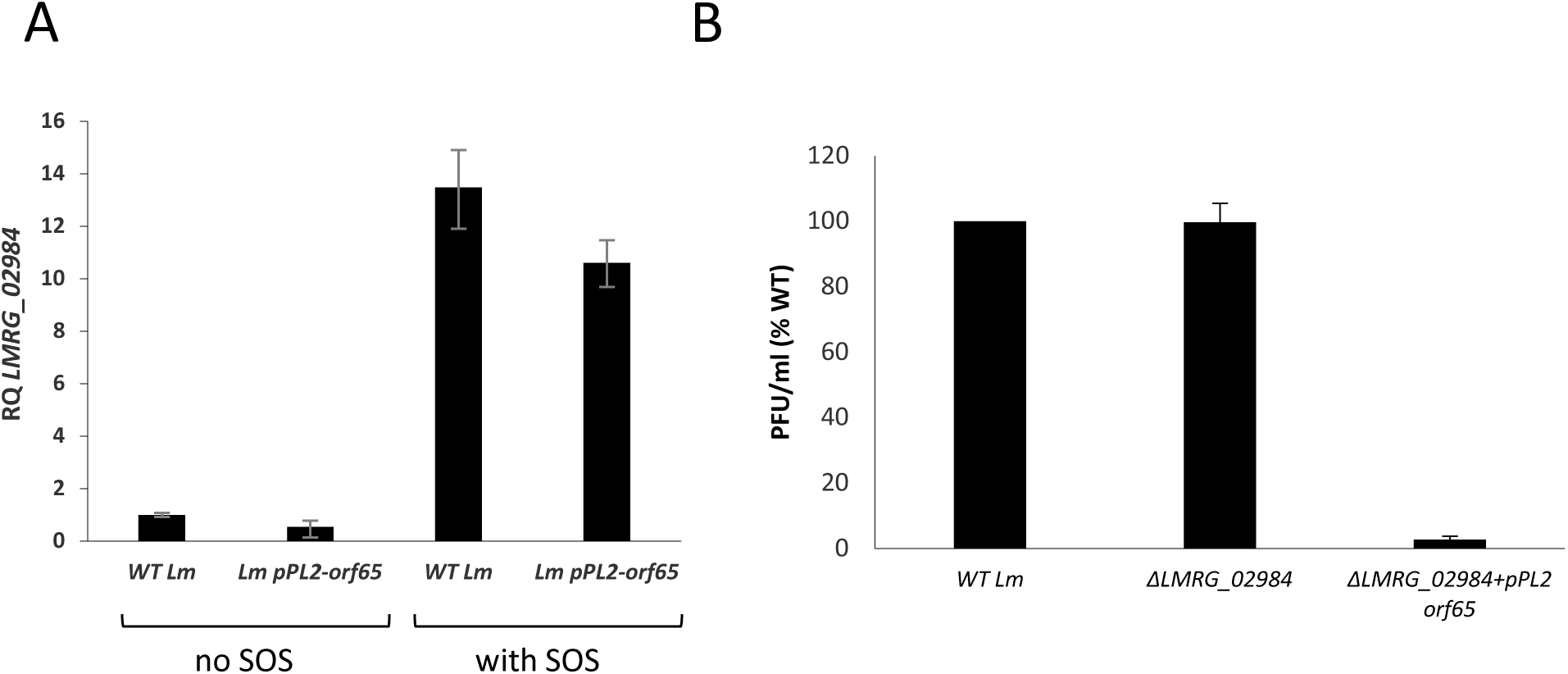
*orf65* does not act as anti-sense RNA of the *LMRG_02984* gene. **A.** Strand-specific RT-qPCR analysis of *LMRG_02984* transcription levels in WT *Lm* and *Lm pPL2-orf65* strains. The bacteria were grown in BHI medium at 30°C and subjected or not to UV irradiation. Analysis was performed 4 h post-UV irradiation. The results are presented as RQ, relative to the levels in WT bacteria. The experiment was performed three times. The figure shows a representative assay. Additional biological repeats can be found in the **Data S1 file**. **B.** A plaque-forming assay of virions obtained from WT *Lm,* Δ*LMRG_02984* and Δ*LMRG_02984 pPL2-orf65* strains upon MC treatment. Virions were isolated from bacterial lysates 6 h post-MC treatment and tested on the Mack861 indicator strain for plaque-forming units (PFU). Error bars represent the standard deviation of three independent experiments.

**Figure S5.**
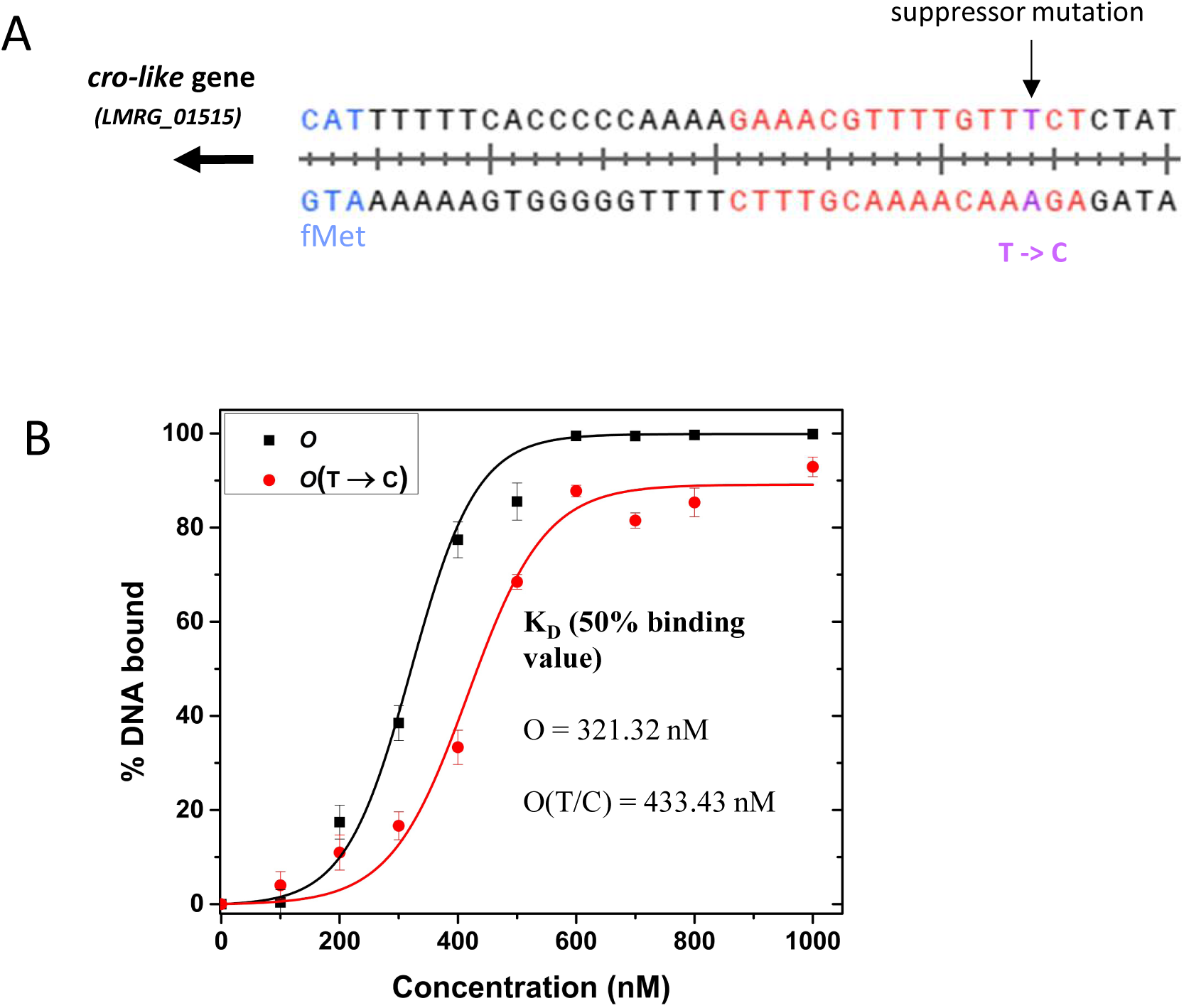
Cro binds the mutated operator with lower affinity. **A.** The region upstream of the *cro-like* gene is shown, where the T-to-C mutation was identified (marked in purple). The putative operator site of *cro* is shown, comprising 16 nucleotides (marked in red). **B.** EMSA analysis of purified Cro proteins binding to DNA probes containing the *cro* operator site with or without the T-to-C mutation. The experiment was performed in at least three independent biological repeats, and the average result is presented. Error bars represent standard deviation. Fitting was performed using Origin.

**Figure S6.**
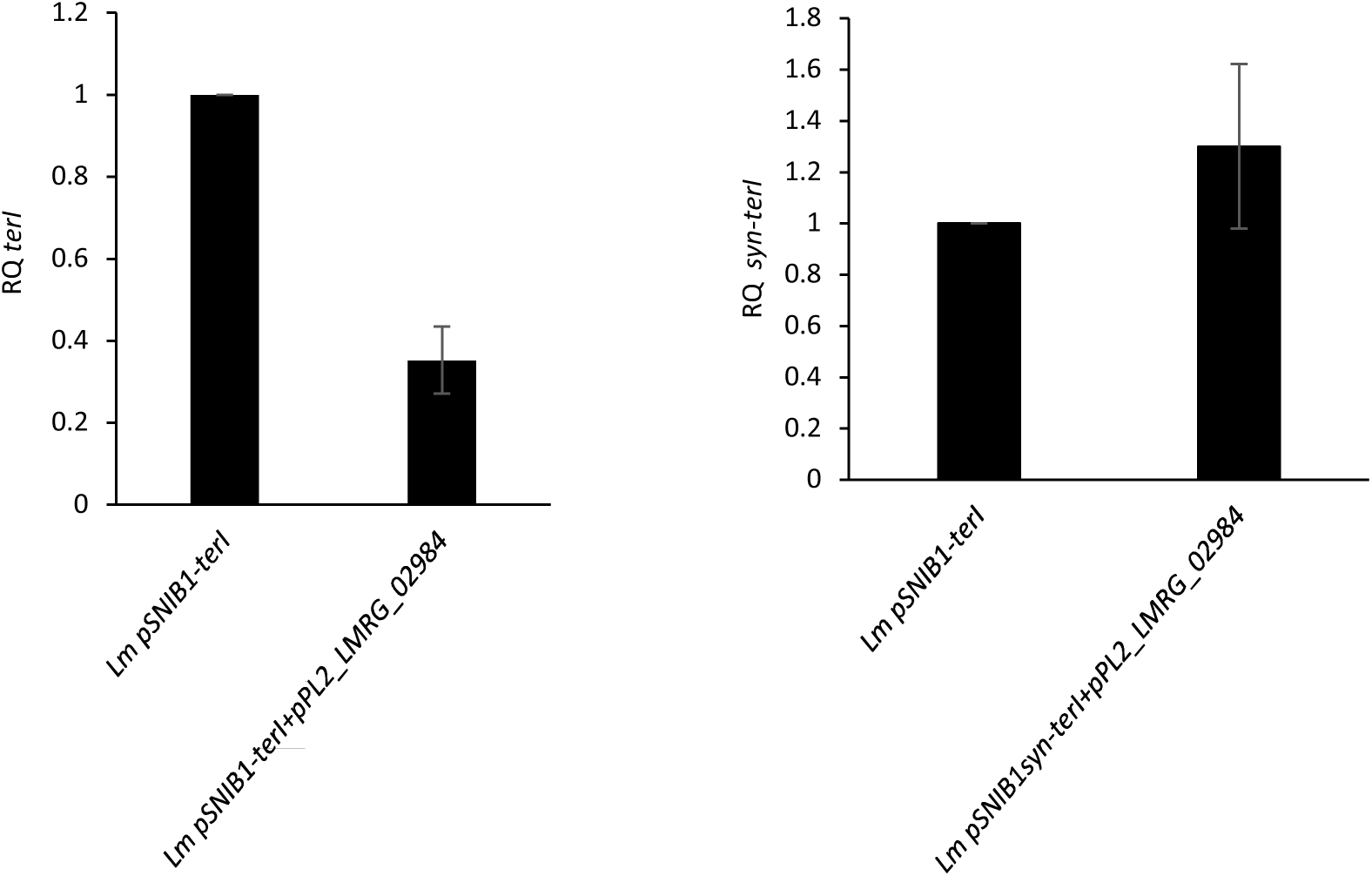
*LMRG_02984* ectopic expression does not silence transcription of the synthetic *terI* gene. Strand-specific RT-qPCR analysis of *terI* transcription in *Lm pSNIB1-terI* (left panel) and *Lm* bacteria expressing the synthetic *terI* gene *(Lm pSNIB1syn-terI,* right panel) in the presence or absence of *LMRG_02984* expression from the pPL2 plasmid. The bacteria were grown in BHI medium for 3 h at 30°C. The results are presented as RQ, relative to their levels in *Lm pSNIB1-terI* and *Lm pSNIB1syn-terI* bacteria, respectively. Error bars represent the standard deviation of three independent experiments.

**Figure S7.**
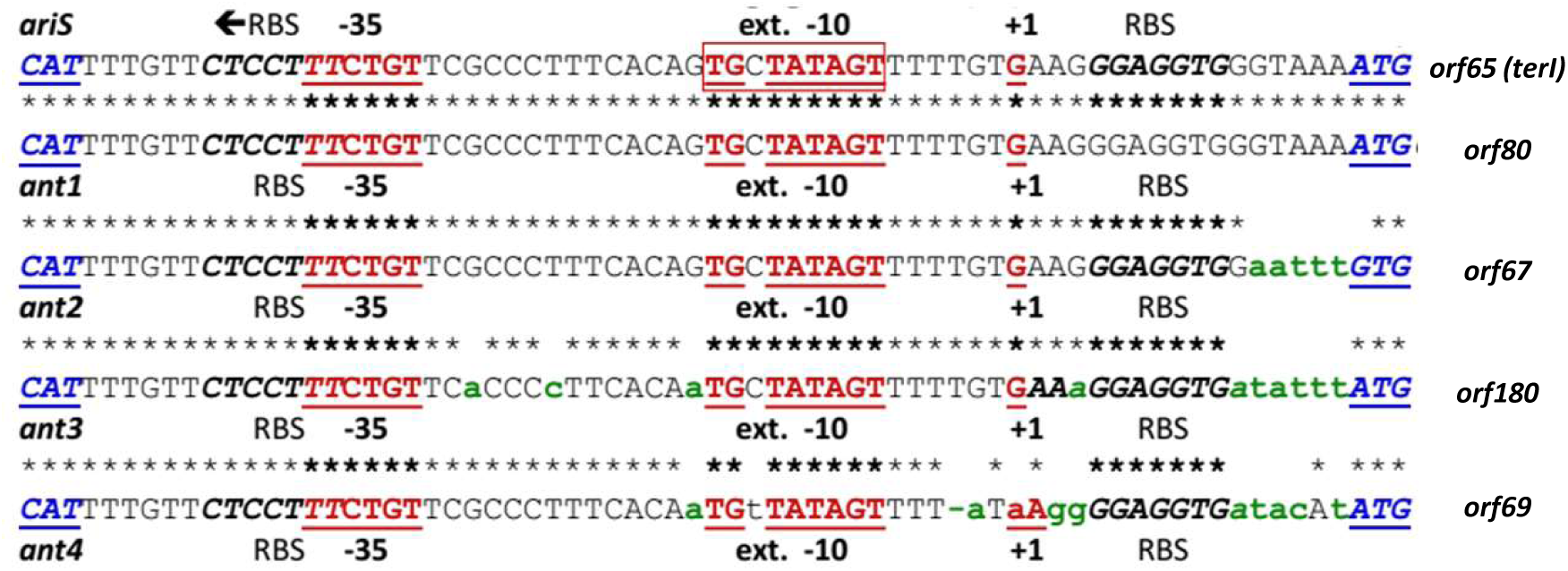
The promoter of *terI* is conserved in *comK-pro*phages. A comparison of *orf65* (*terI*), *orf180, orf80, orf67* and *orf69* promoter regions, demonstrating high similarity. The promoter of *orf65* was taken from the *comK*-prophage of *Lm* strain 10403S, the promoter of *orf180* was taken from the *comK*-prophage of *Lm* strain L1846, the promoter of *orf80* was taken from the *comK*-prophage of *Lm* strain 10-092876-0168, the promoter of *orf69* was taken from the *comK*-prophage of *Lm* strain ATCC 51779 and the promoter of *orf69* was taken from the *comK*-prophage of *Lm* strain PSU-VKH-LP019.

